# Intra-African Geographic Domain Shift in Wildlife Camera Trap Species Classification: A Comparative Study of Supervised and Zero-Shot Foundation Models

**DOI:** 10.64898/2026.06.24.734283

**Authors:** Nanda Nanduri, Jesutomiwo Ogundare, George Anderson

## Abstract

Camera trap networks such as Snapshot Safari have generated millions of labelled wildlife images across Africa, enabling the training of deep learning models for automated species classification. However, deploying models trained in one African region to another remains poorly understood. To the best of our knowledge, this study presents the first systematic evaluation of geographic domain shift within the African continent for wildlife camera trap species classification, using the Machine Learning sub-field of Artificial Intelligence. We use three model architectures, each interacting with Snapshot Serengeti in a different way: BEiTV2 is fine-tuned on Serengeti images as a supervised baseline; DINOv2 with FAISS uses Serengeti images as a retrieval index without any weight updates; and BioCLIP is a true zero-shot foundation model that receives no Serengeti training data at all. All three are then evaluated on two Southern African test sets—Snapshot Kgalagadi and Snapshot Kruger —as well as on locally collected wildlife photographs from Botswana. We conduct eight experiments covering in-domain baselines, cross-dataset transfer, data scaling, MegaDetector preprocessing, grayscale vs. colour image conditions, and per-species transfer analysis. This work provides the first empirical characterisation of intra-African domain shift across both supervised and zero-shot architectures, and offers practical guidance for conservation AI practitioners who need to deploy models across the diverse ecosystems of Southern Africa without collecting new labelled data.

## Introduction

Camera traps have become one of the most valuable tools available to wildlife ecologists and conservation managers across Africa. These motion-activated devices operate continuously in the field without requiring human presence, capturing images of animals across large areas and over extended periods of time (Burton et al., 2015; Tuia et al., 2022). Networks such as Snapshot Safari have deployed hundreds of cameras across multiple African protected areas, from Tanzania in the east to Botswana and South Africa in the south, all following the same standardised protocols to allow meaningful comparisons between sites (Pardo et al., 2021). The Snapshot Serengeti project had accumulated 1.2 million image sets across its first three seasons of operation by 2013 (Swanson et al., 2015), and has since grown to over seven million images across eleven seasons. However, the sheer scale of this data has created a serious practical bottleneck. Manually reviewing and labelling millions of images requires enormous amounts of expert time and is neither sustainable nor scalable as camera trap networks continue to grow (Norouzzadeh et al., 2018; Vélez et al., 2023).

Advances in deep learning (a sub-field of Machine Learning (ML), which is itself a sub-field of Artificial Intelligence (AI)), particularly Convolutional Neural Networks (CNNs), have offered a promising solution to this problem. Several studies have demonstrated that CNNs can classify wildlife species from camera trap images with impressive accuracy when the model is trained and tested within the same geographic area (Gomez Villa et al., 2017; Norouzzadeh et al., 2018; Tabak et al., 2019). These results have encouraged conservation organisations to adopt AI-based classification tools as a way to reduce the burden of manual image review. However, a critical limitation of supervised deep learning approaches is their sensitivity to changes in the visual distribution of images across different environments. When a model is deployed at a new location that was not represented in its training data, performance can degrade dramatically, a phenomenon known as geographic domain shift (Beery et al., 2018). This occurs because models trained in one region tend to rely on background features, lighting conditions, and habitat textures that are specific to that location, rather than on the visual appearance of the animals themselves. As a result, a model that performs well in East Africa may fail when applied to a park in Southern Africa with a different landscape and species assemblage. Santamaria et al. (2026) illustrated the severity of this problem by showing that a vision-language model trained on African camera trap data suffered a top-1 accuracy collapse from 84.77% to just 16.17% when evaluated on data from a different continent.

Despite the growing body of research on domain shift in wildlife AI, the problem has predominantly been examined through the lens of localized, single-region studies in North America (Beery et al., 2018) or large inter-continental transfers (Santamaria et al., 2026). Studies that have applied foundation models to African camera trap data, such as WildCLIP (Gabeff et al., 2024), have evaluated their models exclusively within the same dataset they trained on, leaving cross-regional generalisation entirely untested. Geographic domain shift within the African continent, where ecosystems and species assemblages differ meaningfully between regions but share a degree of ecological continuity, has received virtually no systematic attention. This is a significant gap in the literature. A review of ML studies applied to wildlife imagery by Nakagawa et al. (2023), published in Peer Community Journal, found that most published research originates from a small number of countries and that geographic diversity in study locations remains severely limited. The African continent, despite hosting some of the world’s most biodiverse and conservation-critical ecosystems, is underrepresented in terms of rigorous cross-regional AI evaluation. This is particularly relevant for Southern African countries like Botswana, where conservation practitioners working in environments such as the Kgalagadi Transfrontier Park and the Kalahari basin need AI tools that are reliable and generalisable in their specific ecological context, not just in East Africa where most training data originates.

The emergence of large-scale multimodal language models and vision foundation models offers a potential alternative to location-specific supervised training. Models such as BioCLIP, DI-NOv2, and BEiTV2 are pretrained on vast and diverse datasets and can perform species recognition without requiring any labelled images from the target deployment region (Fabian et al., 2023; Vyskočil and Picek, 2025). Zero-shot and retrieval-based approaches built on these foundation models are particularly attractive in data-scarce conservation contexts, where collecting and labelling enough images to train a conventional CNN from scratch may not be feasible. Whether these models maintain reliable performance under intra-African geographic domain shift, where differences in background, vegetation, lighting, and species composition are ecologically meaningful but less extreme than inter-continental comparisons, has not yet been systematically evaluated. This represents a critical gap in the literature, as conservation practitioners across Southern Africa increasingly require AI tools that generalise reliably across ecologically distinct but geographically connected African ecosystems.

In this paper, we present the first systematic study of geographic domain shift within the African continent for wildlife camera trap species classification, using the ML sub-field of AI. To the best of our knowledge, no prior work has compared supervised transformer models against zero-shot foundation models specifically under intra-African geographic domain shift conditions, nor used Snapshot Kgalagadi and Snapshot Kruger as dedicated cross-regional test sets for this purpose. BEiTV2 is fine-tuned on a stratified subset of 46,471 images from Snapshot Serengeti, a large and well-established East African dataset from Tanzania. The size of this subset is selected on empirical grounds: Schneider et al. (2020) demonstrated that 1,000 or more labelled images per species are sufficient to achieve stable per-class recall above 97%, and our subset provides approximately 3,000 images per species for most of the 17 retained classes, comfortably above this threshold. DINOv2 with FAISS uses Serengeti images as a retrieval index without any weight updates, and BioCLIP operates in a fully zero-shot manner with no exposure to Serengeti training images whatsoever. All three models are then evaluated on two Southern African test sets, Snapshot Kgalagadi from Botswana and Snapshot Kruger from South Africa, as well as on locally collected wildlife photographs from Botswana that were not captured using camera traps.

*Contributions.* The main contributions of this paper are as follows:

- We provide the first systematic empirical evaluation of intra-African geographic domain shift for wildlife camera trap species classification, training on East African data and evaluating on two Southern African datasets.
- We compare three fundamentally different model architectures — BEiTV2 (supervised fine-tuned transformer), DINOv2 with FAISS (retrieval-based zero-shot), and BioCLIP (true zero-shot foundation model) — under the same domain shift conditions, providing the first direct comparison of these approaches in an African cross-regional setting.
- We introduce Snapshot Kgalagadi and Snapshot Kruger as dedicated test sets for cross-regional AI evaluation, and contribute a novel locally collected wildlife photography dataset from Botswana.
- We conduct eight experiments covering in-domain baselines, cross-dataset transfer, data scaling, MegaDetector preprocessing, grayscale versus colour conditions, and per-species transfer analysis, producing practical guidance for conservation AI practitioners working across Southern African ecosystems.

The remainder of this paper is organised as follows. Section 2 reviews related work on ML for camera trap image classification, multimodal foundation models for wildlife recognition, and geographic domain shift in wildlife AI. Section 3 describes the methodology, including the datasets used, model architectures, experimental design, evaluation metrics, and hyperparameter configurations. Section 4 presents the results of all eight experiments. Section 5 discusses the findings in the context of geographic domain shift, compares results against prior work, and outlines the limitations of this study. Section 6 concludes the paper with practical recommendations for conservation AI practitioners working across Southern African ecosystems and directions for future research.

## Related Work

### Machine Learning and CNNs for Camera Trap Image Classification

The application of machine learning to automate the processing of camera trap images has grown rapidly over the past decade, driven by the increasing availability of large annotated datasets and advances in deep learning architectures. Early approaches to automated wildlife classification relied on hand-crafted visual features and traditional machine learning algorithms, achieving limited accuracy on small datasets. For example, Yu et al. (2013) used sparse coding with a linear Support Vector Machine to classify 18 species from 7,196 images, reaching a top-1 accuracy of 82%, but only after manually cropping animals from images. Gomez Villa et al. (2017) demonstrated that very deep convolutional neural networks could substantially outperform these earlier methods on the Snapshot Serengeti dataset, achieving 88.9% top-1 accuracy on 26 species, marking a clear turning point in automated wildlife classification.

Norouzzadeh et al. (2018) provided one of the most influential early demonstrations that CNNs could reliably classify wildlife from camera trap images at scale. Working with the Snapshot Serengeti dataset—the same dataset we use for training in this study—they trained several deep neural network architectures and showed that the models could identify 48 species achieving a top-1 accuracy of 93.8% on the expert-labelled test set, match the performance of crowdsourced human volunteers, and automate the labelling of over 99% of images. This was a landmark result that demonstrated the practical potential of deep learning for large-scale wildlife monitoring. Building on this foundation, Tabak et al. (2019) developed a CNN-based classification system trained on over three million images from five locations across the United States, achieving a top-1 accuracy of 98% on an independent test subset drawn from the same geographic region as the training data, and demonstrating that the approach could be packaged into an accessible software tool for ecologists. These results were highly influential in driving the adoption of CNN-based tools by conservation practitioners.

Subsequent work expanded the scope of CNN-based wildlife classification in several important directions. Willi et al. (2019) showed that applying transfer learning from CNNs pretrained on large camera trap datasets was particularly effective for smaller projects where labelled images were limited, raising classification accuracy by up to 10.3% in those cases and substantially reducing the time needed for model training. They also demonstrated that combining model predictions with citizen science classifications could reduce human labelling effort by 43% while maintaining overall accuracy. Schneider et al. (2020) specifically investigated deep learning performance on a modestly sized dataset of 47,279 images from 36 locations in Canada, demonstrating that classification accuracy above 95% can be achieved even with limited training data when using transfer learning and image augmentation, and providing a practical guideline that 1,000 or more labeled images per species are required to achieve a stable per-class recall of 0.971 (97.1%). Vélez et al. (2023) conducted a systematic evaluation of four widely used AI platforms for camera trap processing—MegaDetector, Conservation AI, MLWIC2, and Wildlife Insights— testing them on datasets from four geographically distinct regions: North America (Montana), South America (Orinoquía Camera Traps), Africa (Snapshot Kgalagadi), and Asia (SWG Camera Traps). They showed that species classification performance dropped substantially when models were applied to datasets outside their training distribution, with per-species F1 scores falling below 0.5 for the majority of taxa on the Kgalagadi dataset and some common species scoring near zero. This finding is directly relevant to our study as it demonstrates that even well-resourced and widely adopted tools struggle with geographic domain shift. More recently, Mulero-Pázmány et al. (2025) proposed a two-stage deep learning framework using MegaDetectorV5 as a backbone and trained specialist models for different animal groups, achieving an F1 score of 96.2% on a dataset of 1.3 million images but again within a single geographic region.

Despite the impressive performance reported in many of these studies, they share a common limitation that motivates our work directly. Nearly all supervised approaches in this area are evaluated within the same geographic region where the training data was collected, and performance under genuine cross-regional conditions is rarely examined. Where cross-dataset evaluation has been attempted, the results reveal that accuracy degrades significantly (Beery et al., 2018; Vélez et al., 2023). Our study differs from all of the above in that we deliberately use East African training data and evaluate on two Southern African test sets, treating geographic generalisation as the primary object of study rather than a secondary observation. Furthermore, while previous work has relied primarily on CNN and ResNet-style architectures, we evaluate transformer-based and foundation model approaches to assess whether these newer architectures offer genuinely better generalisation under domain shift conditions.

### Multimodal Language Models and Zero-Shot Approaches for Wildlife

While supervised CNN approaches have dominated camera trap research, a growing body of work has explored whether large-scale foundation models can classify wildlife species without requiring labelled training data from the target deployment region. These zero-shot and retrieval-based approaches are particularly appealing for conservation contexts where annotated datasets are unavailable, expensive to produce, or rapidly outdated as cameras are deployed in new locations.

Radford et al. (2021) introduced CLIP, a vision-language model pretrained on hundreds of millions of image-text pairs collected from the internet. CLIP demonstrated that it was possible to classify images of unseen categories simply by comparing image features against natural language descriptions of those categories, without any task-specific training. This established the foundation for applying vision-language models to wildlife classification in a zero-shot manner. Building on this, Gabeff et al. (2024) created WildCLIP by fine-tuning CLIP specifically on camera trap images from the Snapshot Serengeti dataset and adding an adapter module that allows the model to learn new ecological attributes from a small number of examples. WildCLIP demonstrated that vision-language models could retrieve camera trap images according to complex ecological descriptions such as species appearance, behaviour, and scene attributes. Importantly, WildCLIP was evaluated exclusively within the Snapshot Serengeti dataset, which is our training dataset in this study, and its cross-regional generalisation was never tested.

Fabian et al. (2023) proposed a different zero-shot pipeline called WildMatch that requires no location-specific training data at all. Their system used a vision-language model to generate detailed natural language descriptions of animals visible in camera trap images and then matched those descriptions against a knowledge base of species descriptions extracted from Wikipedia. Without any training on local data, WildMatch achieved a top-1 accuracy of 62.3% on a Colombian camera trap dataset and substantially outperformed standard CLIP baselines, which reached only 41.7% on the same data. This demonstrated for the first time that zero-shot approaches based on large language models could be competitive with supervised methods in genuinely novel deployment environments where no prior training data exists.

Vyskočil and Picek (2025) provided a comprehensive comparison of transformer-based and foundation model architectures for camera trap classification, testing BEiTV2, DINOv2 with FAISS-based retrieval, BioCLIP, BLIP, and ChatGPT across three datasets from New Zealand, the United States, and Central Europe. Their experiments showed that BEiTV2 achieved top-1 accuracy of 84.2%, 92.2%, and 96.6% on the CCT20, CEF22, and WCT datasets respectively, significantly outperforming traditional CNN baselines, particularly for rare species. When combined with MegaDetector preprocessing, the relative top-1 accuracy error of a single BEiTV2 classifier was further reduced by between 42% and 75% depending on the dataset. They also showed that DINOv2 with FAISS retrieval achieved competitive zero-shot performance without any model training, and that this retrieval approach was especially useful for small datasets where supervised training was difficult. These findings directly motivated our choice of BEiTV2 and DINOv2 with FAISS as two of the three model architectures we evaluate in this study.

On a broader scale, Nakagawa et al. (2023), in a systematic review of over two hundred papers on ML for wildlife imagery published in Peer Community Journal, found that the field is moving rapidly towards deep learning approaches but that geographic diversity in study locations remains very limited, with most published research originating from India, China, Australia, and the United States, and Africa underrepresented despite hosting exceptional wildlife diversity. The third model we evaluate is BioCLIP, introduced by Stevens et al. (2024), which was pretrained specifically on biological imagery using taxonomic information from the tree of life, making it the most biologically informed of the foundation models we test.

Our work builds directly on this body of research but addresses a gap that none of the above studies examined. Existing applications of zero-shot and foundation models to camera trap data have either been evaluated on a single dataset (Gabeff et al., 2024) or heavily relied on inter-continental datasets spanning different continents (Fabian et al., 2023; Vyskočil and Picek, 2025). Whether supervised transformers and zero-shot foundation models differ meaningfully in their robustness to intra-African geographic domain shift has not been examined. By evaluating BEiTV2, DINOv2 with FAISS, and BioCLIP on Southern African test sets—where only BEiTV2 uses East African training data and the zero-shot models require no local data at all—our study provides the first assessment of the practical trade-offs between supervised and zero-shot approaches for conservation practitioners deploying AI across ecologically distinct but geographically connected African ecosystems.

### Domain Shift in Wildlife Camera Trap AI

The problem of domain shift, where a model trained on data from one distribution performs poorly when tested on data from a different distribution, has been recognised as one of the most significant practical barriers to deploying wildlife AI systems in real conservation settings. In the context of camera trap imagery, domain shift typically arises when models are applied to new camera locations with different backgrounds, vegetation, lighting conditions, or species assemblages that were not represented during training.

Beery et al. (2018) established the foundational framework for understanding domain shift in camera trap AI through their Terra Incognita dataset and evaluation. They demonstrated that models achieving strong performance on seen camera locations experienced severe degradation when tested on cameras from unseen locations within the same broad geographic region. Their analysis showed that this failure was primarily driven by the tendency of neural networks to learn location-specific background features rather than the actual appearance of the animals, a finding that has since been confirmed by multiple subsequent studies. This paper remains the primary reference point for understanding geographic generalisation failure in wildlife AI and directly motivates the experimental design of our study.

Subsequent work attempted to quantify and address domain shift through various approaches. The iWildCam challenge series provided benchmark datasets specifically designed to test cross-location generalisation, helping the research community develop a shared evaluation standard for out-of-distribution performance in wildlife classification (Beery et al., 2019a). The 2019 edition trained models on data from the American Southwest and tested on the American Northwest, explicitly enforcing geographic disjointness to simulate real-world deployment conditions. Vélez et al. (2023) provided empirical evidence of domain shift in practice by applying widely used AI platforms to the Snapshot Kgalagadi dataset, which forms one of our two primary test sets. They showed that species classification performance dropped substantially when models were applied to datasets outside their training distribution, testing across four geographically distinct regions — North America (Montana), South America (Orinoquía), Africa (Snapshot Kgalagadi), and Asia (SWG Camera Traps) — with per-species F1 scores falling below 0.5 for the majority of taxa on the Kgalagadi dataset and some common species scoring near zero. This finding is directly relevant to our study because it demonstrates that the challenge we are investigating is real and practically significant even for well-resourced, widely adopted AI tools.

More recently, domain shift research has moved towards foundation models as a potential solution. Gabeff et al. (2024) showed through WildCLIP that fine-tuning a vision-language model on Serengeti camera trap images could improve retrieval performance, but their evaluation remained within the Serengeti dataset and did not test cross-regional transfer to other African ecosystems. Santamaria et al. (2026) made the most direct contribution to this problem with their WildIng model, which was explicitly designed to be robust to geographical domain shift. They trained on Snapshot Serengeti and tested on the Terra Incognita dataset from the United States, documenting a top-1 accuracy collapse from 84.77% to 16.17% for a standard vision-language model baseline. Their WildIng approach, which combined visual features with text descriptions of species appearance, recovered approximately 30% of this lost performance by learning geographically invariant representations. While this is an important step forward, their evaluation was limited to an Africa-to-America transfer scenario where the domain gap is very large and the species assemblages are almost entirely different. Vyskočil and Picek (2025) also touched on location generalisation and found that transformer-based models overfit less to specific camera locations than CNN-based models, with the performance gap between seen and unseen locations being consistently smaller for transformer architectures.

Domain shift research within the African continent specifically, between ecosystems that share some species but differ significantly in habitat, vegetation, and environmental conditions, has not been addressed in the literature. Studies using African camera trap data have almost exclusively evaluated their models on the same dataset they trained on (Gabeff et al., 2024; Norouzzadeh et al., 2018; Willi et al., 2019), or have applied pre-existing models to African data without training new ones specifically for cross-regional evaluation (Vélez et al., 2023). The question of how well a model trained on East African savanna data generalises to the arid Kalahari ecosystem of Southern Botswana, or to the mixed woodland habitats of Kruger National Park in South Africa, has not been studied.

Our paper directly fills this gap. Unlike Santamaria et al. (2026) who crossed continents, and unlike Beery et al. (2018) who worked within North America, we examine domain shift between ecologically distinct but geographically connected African ecosystems. This represents a more realistic and practically relevant evaluation scenario for conservation AI deployment across the African continent, where practitioners increasingly want to leverage large East African training datasets to build tools that work reliably in Southern Africa. This is the first study to systematically evaluate intra-African geographic domain shift using both supervised and zero-shot AI models, and the first to use Snapshot Kgalagadi and Snapshot Kruger as dedicated test sets for this purpose.

## Methodology

### Datasets

This study uses four datasets representing distinct geographic regions and image collection contexts across the African continent. Three of these datasets are publicly available through the Labeled Information Library of Alexandria for Biology and Conservation (LILA BC), each collected as part of the Snapshot Safari network using standardised camera trap protocols that allow direct cross-site comparisons (Pardo et al., 2021). The fourth dataset consists of locally collected wildlife photographs from Botswana.

#### Snapshot Serengeti (Training Set)

Snapshot Serengeti is the training dataset for this study and serves as the source domain in all cross-regional experiments. It covers Serengeti National Park in Tanzania, a large East African savanna ecosystem characterised by open grasslands, dense woodlands, and one of the highest concentrations of large mammals on earth. The dataset contains approximately 7.1 million images from 2.65 million sequences, where a sequence refers to a burst of one to three photographs triggered by a single motion detection event at a camera station, collected across eleven seasons of continuous camera trap operation using a grid of 225 cameras covering 1,125 square kilometres (Swanson et al., 2015). Labels are provided for 61 species categories, with the most frequently occurring species being wildebeest, zebra, and Thomson’s gazelle. Approximately 76% of all images are labelled as empty. The dataset is released under the Community Data License Agreement permissive variant and is freely available through LILA BC.

#### Snapshot Kgalagadi (Test Set 1)

Snapshot Kgalagadi covers the Kgalagadi Transfrontier Park, a large arid savanna ecosystem spanning the border between Botswana and South Africa in the Kalahari region, with the park’s western boundary bordering Namibia. This park represents a very different ecological environment from Serengeti, with sparse desert vegetation, extreme temperature variation, and a species assemblage dominated by desert-adapted animals. The dataset contains 10,222 images from 3,611 sequences and provides labels for 31 species categories, with the most common being gemsbok/oryx, other birds, and ostrich (LILA BC, 2024a). Approximately 76.14% of images are labelled as empty. This dataset is the primary Southern African test set in our study and has not previously been used as a dedicated test target for cross-regional AI evaluation.

#### Snapshot Kruger (Test Set 2)

Snapshot Kruger covers Kruger National Park in South Africa, one of the most species-rich protected areas remaining on the African continent, established in 1898. The park encompasses mixed woodland and savanna habitats and houses a diverse assemblage of large mammals including the full complement of the so-called Big Five. The dataset contains 10,072 images from 4,747 sequences with labels for 46 species categories, the most common being impala, elephant, and buffalo (LILA BC, 2024b). Approximately 61.60% of images are labelled as empty. Kruger represents an intermediate ecosystem between the open plains of Serengeti and the arid landscape of Kgalagadi, allowing us to examine whether domain shift severity varies with the degree of ecological difference between source and target regions.

#### Local Botswana Images (Test Set 3)

The third test set consists of wildlife photographs collected locally in Botswana by the research team using standard photography equipment rather than camera traps. These images represent an additional and distinct form of domain shift because they differ from the Serengeti training data not only in geographic location and species composition but also in image capture conditions, camera type, resolution, and viewing angle. Including this dataset allows us to assess how well the models generalise to wildlife imagery that departs more substantially from the controlled camera trap format on which all three models were trained or evaluated. This test set is a novel contribution of this study and has not appeared in any prior published work. Species identifications for the baboon images in this dataset were confirmed by a wildlife ecologist at the Okavango Research Institute, University of Botswana.

#### Species Overlap Across Datasets

Species overlap analysis was conducted using the combined all-seasons Snapshot Serengeti metadata spanning Seasons 1 through 11, which contains 1,732,540 wildlife annotations across 50 species categories. Table 1 summarises the total image counts and species categories for each dataset before the application of any filtering threshold. Species names were first standardised to a shared canonical vocabulary to account for naming inconsistencies across datasets. For example, Kgalagadi labels gemsbok as *gemsbokoryx*, Kruger records two jackal species separately as *jackalblackbacked* and *jackalsidestriped*, and Serengeti splits lion into *lionfemale*, *lionmale*, and *lioncub*. These were all mapped to single canonical names (gemsbok, jackal, and lion respectively) before comparison. Non-wildlife categories such as empty frames, human, and unidentified birds were excluded from all overlap calculations. The normalised species lists were then compared programmatically using set intersection to identify shared species and their corresponding image counts across datasets.

**Table 1.** Summary of all four datasets used in this study, showing total images and the number of species categories before filtering.

| Dataset | Total images | Species categories |
| --- | --- | --- |
| Snapshot Serengeti (all seasons) | ≈7.1M | 61 |
| Snapshot Kgalagadi | 10,222 | 31 |
| Snapshot Kruger | 10,072 | 46 |
| Local Botswana | 316 | 6 |

Serengeti and Kgalagadi share 22 species after normalisation, including ostrich, jackal, kori bustard, eland, wildebeest, spotted hyena, cheetah, leopard, lion, hartebeest, caracal, porcupine, honey badger, secretary bird, hare, bat-eared fox, aardvark, steenbok, wildcat, duiker, kudu, and brown hyena. Serengeti and Kruger share a larger set of 33 species, reflecting Kruger’s greater ecological similarity to Serengeti, including baboon, buffalo, elephant, giraffe, zebra, impala, hippo, warthog, waterbuck, dik-dik, jackal, hare, spotted hyena, wildebeest, reedbuck, serval, civet, genet, vervet monkey, aardwolf, rhinoceros, and others. For the local Botswana test set, 7 species overlap with the Serengeti training set: elephant (177 images), baboon (52), impala (34), hippo (23), giraffe (19), buffalo (11) and lions (ommitted due to insufficient images), totalling 316 photographs. It should be noted that the local Botswana images are standard wildlife photographs rather than camera trap sequences, and the same individual animal may appear across multiple images.

Image counts for shared species vary substantially across datasets. For example, Serengeti contains 533,478 wildebeest annotations across all seasons while Kgalagadi contains only 13, and spotted hyena has 21,998 annotations in Serengeti but only 1 in Kgalagadi. Because the focus of this study is geographic domain shift at the ecosystem level rather than rare-species classification per se, we apply a minimum threshold of 9 images per species in the target dataset to ensure that per-class metrics are estimated on a sufficient number of test examples to be interpretable. Species below this threshold are excluded from the cross-domain evaluation, with the understanding that single-image or near-single-image species cannot support reliable per-class precision, recall, or F1 estimates and would otherwise destabilise the macro F1 metric used as the primary evaluation criterion. The threshold of 9 was selected as the smallest count for which a model achieving the worst possible per-class score (zero correct predictions) is clearly distinguishable from one with even modest performance, while retaining as many shared species as possible. After applying this threshold, 5 species were retained for Experiment 2 (Serengeti to Kgalagadi): ostrich (46), jackal (39), kori bustard (34), eland (14), and wildebeest (13), yielding a total of 146 test images. For Experiment 3 (Serengeti to Kruger), 14 species were retained: impala (585), elephant (204), zebra (112), buffalo (96), giraffe (94), spotted hyena (78), hare (68), hippo (53), warthog (46), waterbuck (43), baboon (38), wildebeest (23), dik-dik (19), and jackal (9), yielding a total of 1,468 test images.

#### Image Analysis

Image analysis was conducted across all four datasets to characterise their visual properties and assess potential sources of domain shift beyond species overlap. Resolution analysis was performed because native image resolution varies substantially across camera trap deployments and resizing images to fit model input dimensions causes information loss that differs across datasets (Mulero-Pázmány et al., 2025). Class distribution was examined because the absolute and relative frequency of species in a dataset directly affects classification accuracy, with rare and underrepresented species consistently showing lower recall (Schneider et al., 2020; Willi et al., 2019). The proportion of colour versus grayscale images was analysed separately from day versus night capture proportions because these two properties are not equivalent — not all nighttime captures produce grayscale images, and not all grayscale images are necessarily nighttime captures (Tabak et al., 2019). Day versus night analysis was conducted because classification performance has been shown to differ between daytime and nighttime conditions, with daytime accuracy of 98.2% compared to 96.6% at night (Tabak et al., 2019), and because certain foundation models have been shown to fail entirely on infrared images (Vyskočil and Picek, 2025). Brightness statistics were computed to characterise the overall illumination profile of each dataset, as lighting conditions have been identified as a factor affecting automated species recognition performance in camera trap applications (Schneider et al., 2020). Representative sample image grids were produced for each dataset to provide qualitative visual context for the quantitative analyses, following standard practice in camera trap classification studies (Mulero-Pázmány et al., 2025; Norouzzadeh et al., 2018; Vyskočil and Picek, 2025).

Prior to image analysis and model evaluation, empty frames containing no animals were identified and excluded from all experiments. Empty images constitute a substantial proportion of each camera trap dataset, representing approximately 76% of Snapshot Serengeti images, 76.14% of Snapshot Kgalagadi images, and 61.60% of Snapshot Kruger images. Including empty frames in species classification evaluation would artificially inflate accuracy by allowing models to trivially reject non-animal images (Vélez et al., 2023). Empty frame removal was performed using the species label metadata provided with each dataset, retaining only images annotated with a wildlife species label. The local Botswana dataset contains no empty frames as all images were deliberately photographed by the research team.

Camera trap images across all three Snapshot Safari datasets also contain text overlays burned directly into the image, including timestamps, camera identification codes, temperature readings, and moon phase indicators. These overlays introduce a source of visual noise that deep learning models may learn to associate with specific camera locations rather than with animal appearance, potentially contributing to location-specific overfitting (Beery et al., 2018). Schneider et al. (2020) noted that camera trap images from real field deployments are inherently messy, containing varied lighting, partial occlusions, and camera-added information that complicate automated classification. No text overlay removal was applied in this study, meaning the models were evaluated under realistic field conditions where such overlays are present. The local Botswana photographs do not contain such overlays as they were captured using standard handheld cameras.

Resolution varies considerably across datasets. Serengeti images have a mean resolution of approximately 2,208*×*1,656 pixels, Kgalagadi images are uniformly captured at 2,592*×*2,000 pixels, and Kruger images are substantially higher resolution at 5,152*×*3,909 pixels. Local Botswana photographs were taken using handheld cameras and vary in resolution depending on the device used. All images are resized to 224*×*224 pixels before being passed to all three models.

DINOv2-Large supports this input resolution via positional-embedding interpolation; this is below the 518*×*518 most commonly used in DINOv2-L benchmarks, and we report the resulting in-domain accuracy as our reference point with the understanding that higher-resolution inference may yield further gains. Differences in native resolution do not directly affect model input but may influence the amount of visual detail retained after downsampling (Mulero-Pázmány et al., 2025). Representative sample images from each dataset are shown in Figures 1, 2, 3, and 4.

**Figure 1.**
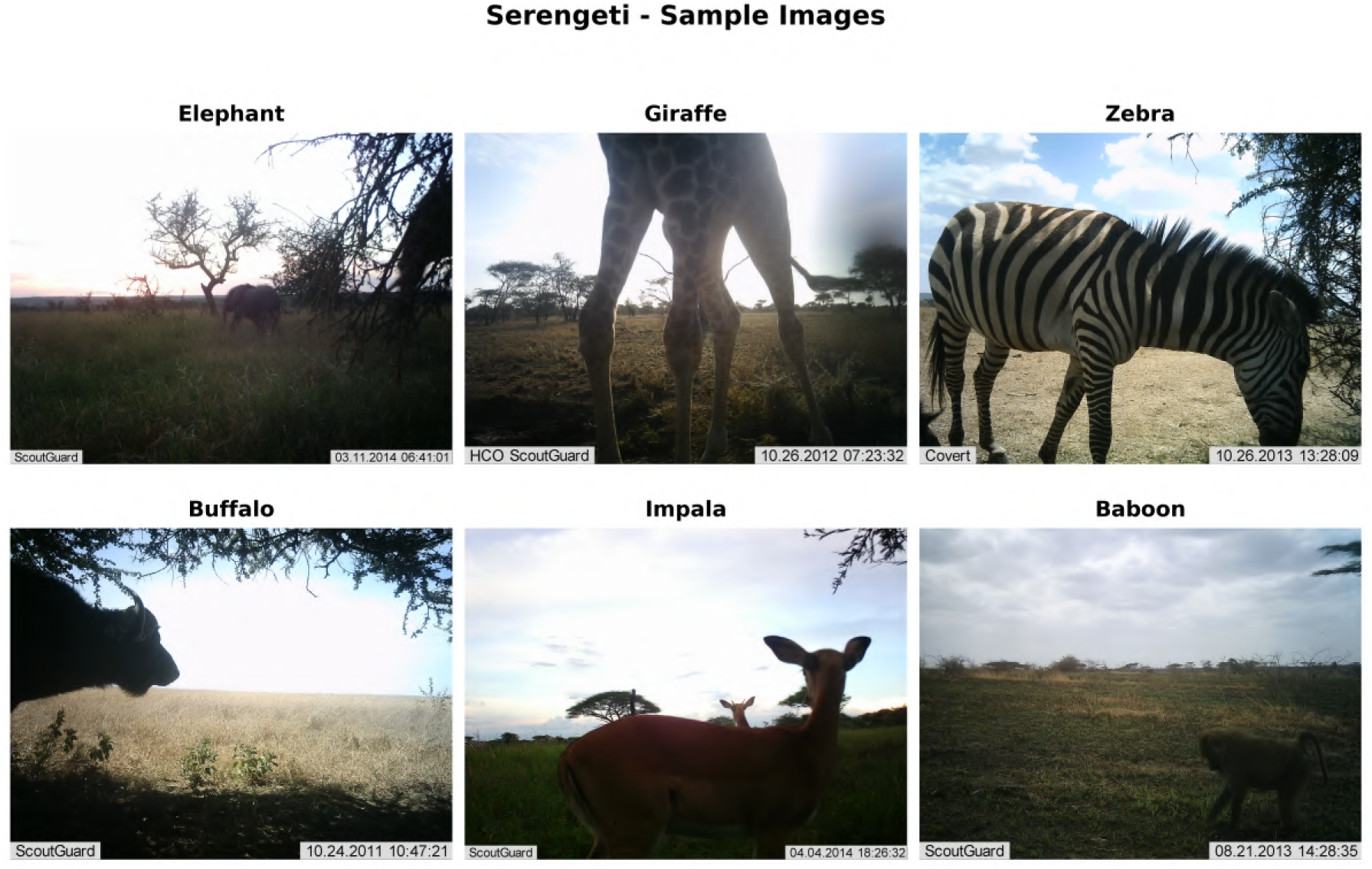
Representative camera trap images from the Snapshot Serengeti training dataset showing six of the 17 training species: elephant, giraffe, zebra, buffalo, impala, and baboon. Images illustrate the open grassland environment and the diversity of lighting and animal positioning typical of East African savanna camera trap deployments.

**Figure 2.**
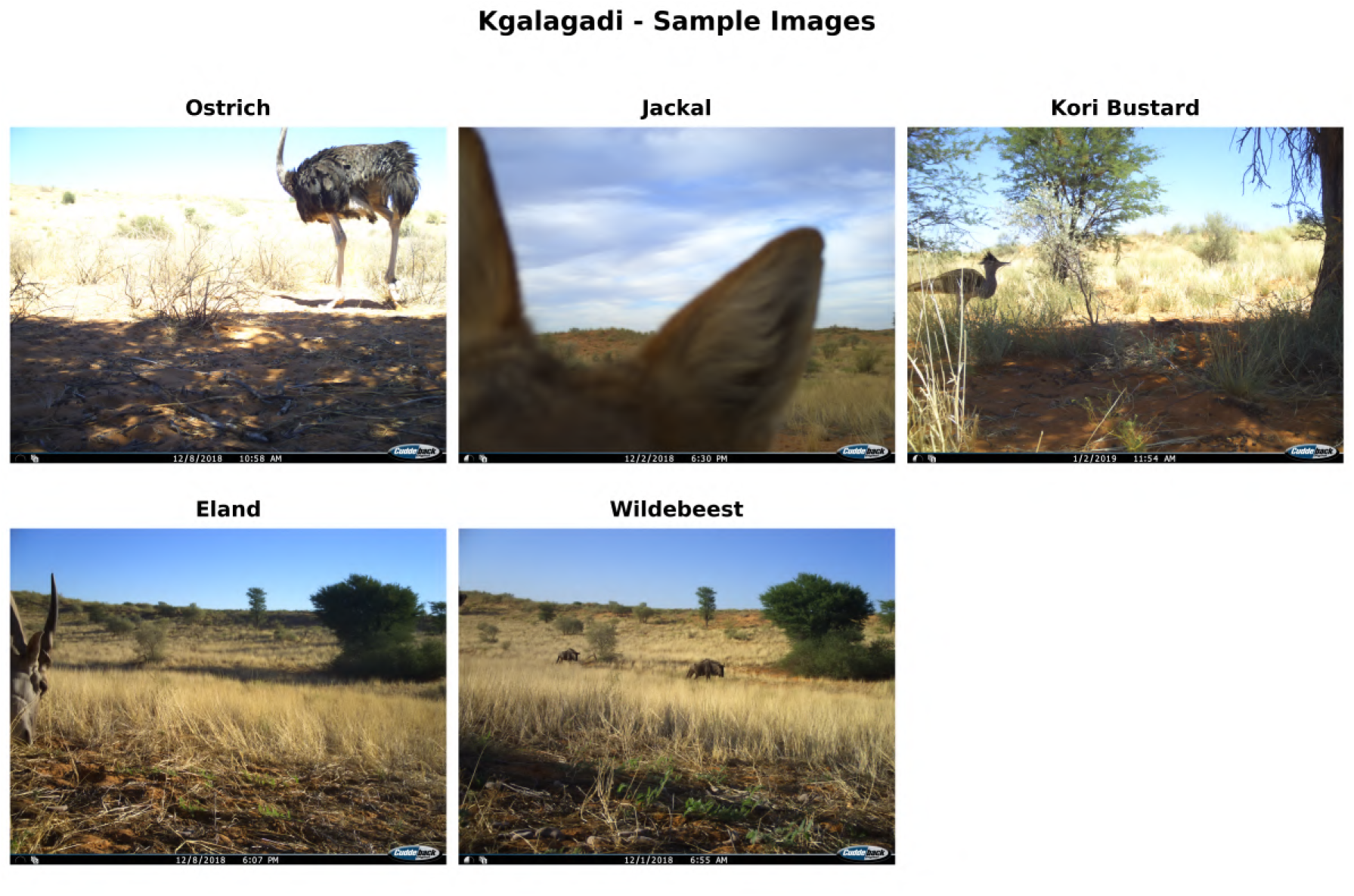
Representative camera trap images from the Snapshot Kgalagadi test set showing all five shared species retained after filtering: ostrich, jackal, kori bustard, eland, and wildebeest. Images illustrate the sparse desert vegetation and arid conditions of the Kalahari ecosystem, which differs substantially from the Serengeti training environment.

**Figure 3.**
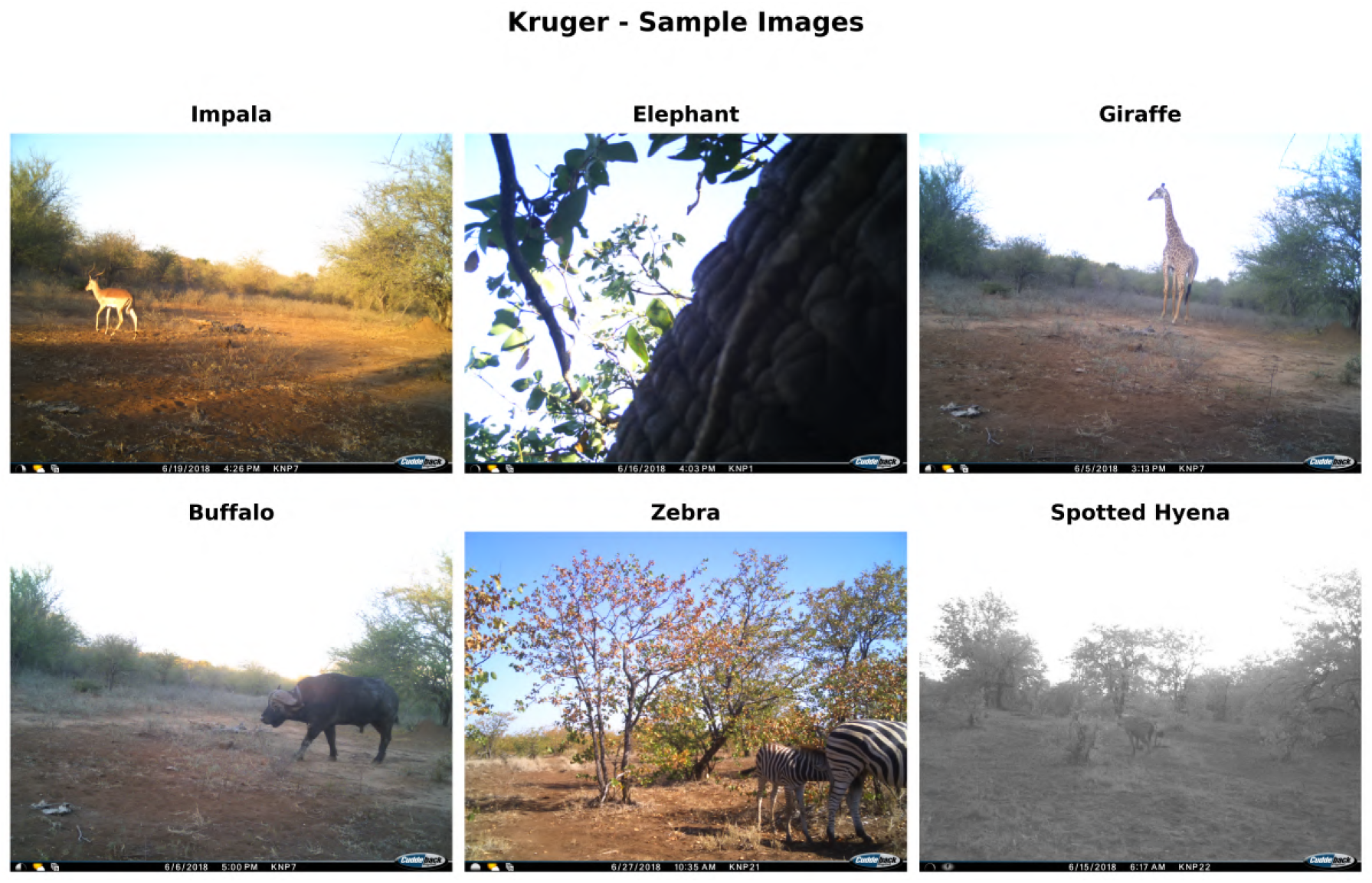
Representative camera trap images from the Snapshot Kruger test set showing six of the 14 shared species: impala, elephant, giraffe, buffalo, zebra, and spotted hyena. Images illustrate the mixed woodland savanna habitat of Kruger National Park, including both daytime colour and nighttime infrared captures.

**Figure 4.**
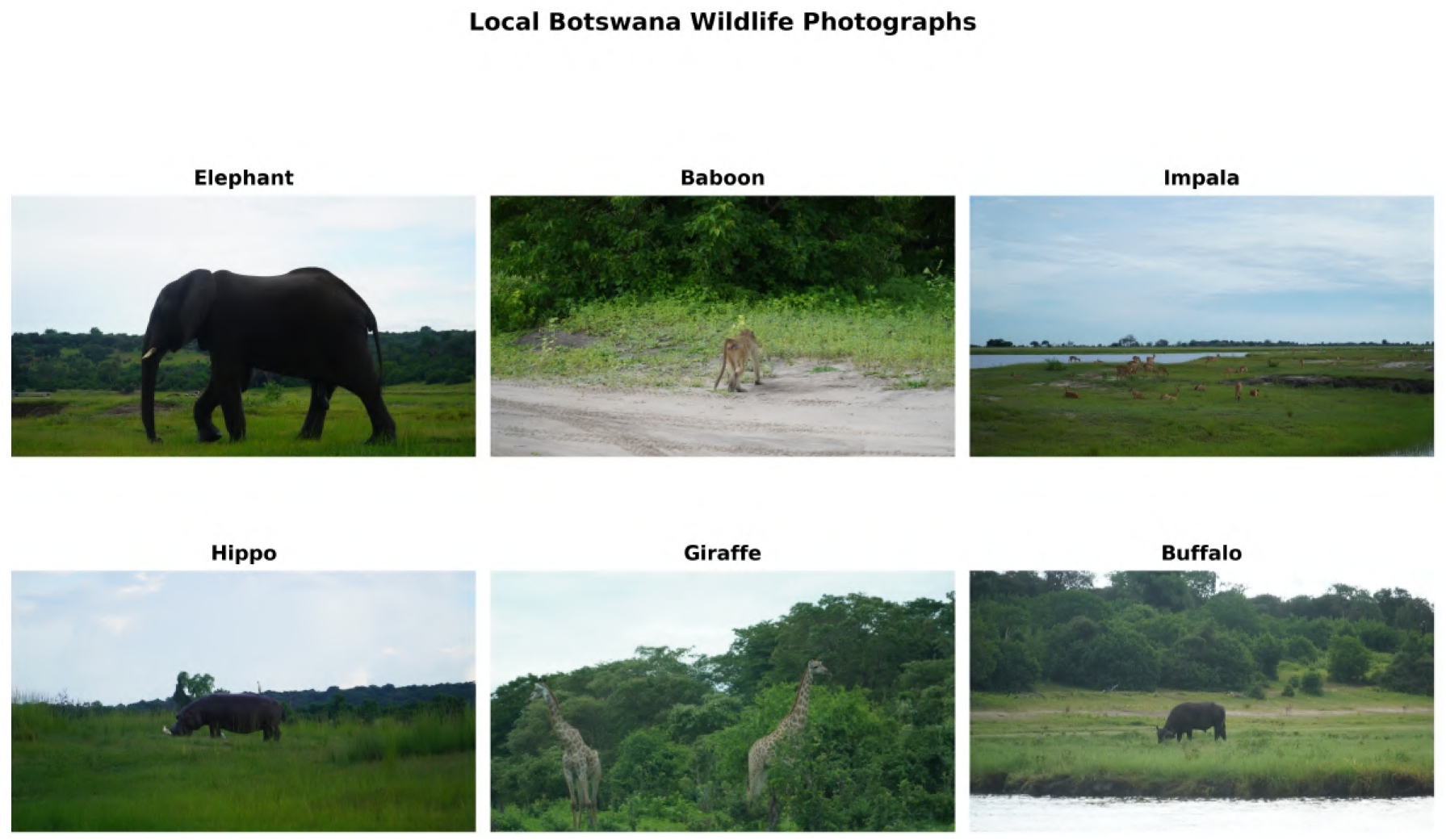
Representative wildlife photographs from the Local Botswana dataset showing all six species: elephant, baboon, impala, hippo, giraffe, and buffalo. Unlike the camera trap images in the other three datasets, these are standard wildlife photographs taken by the research team using handheld cameras during daylight hours, resulting in higher image quality and closer animal proximity than typical camera trap captures.

The proportion of colour versus grayscale images was assessed through pixel-level analysis of 100 randomly sampled images per dataset (Figure 5). An image was classified as grayscale if the standard deviation of the per-pixel channel differences between the red-green and red-blue channel pairs both fell below a threshold of 5, indicating near-equal channel values consistent with infrared capture (Tabak et al., 2019). Serengeti contains 93.0% colour images and 7.0% grayscale images. Kgalagadi contains exclusively colour images at 100.0%. Kruger contains 82.0% colour images and 18.0% grayscale images. All local Botswana photographs are 100% colour, confirmed as all images were collected during daylight hours by the research team.

**Figure 5.**
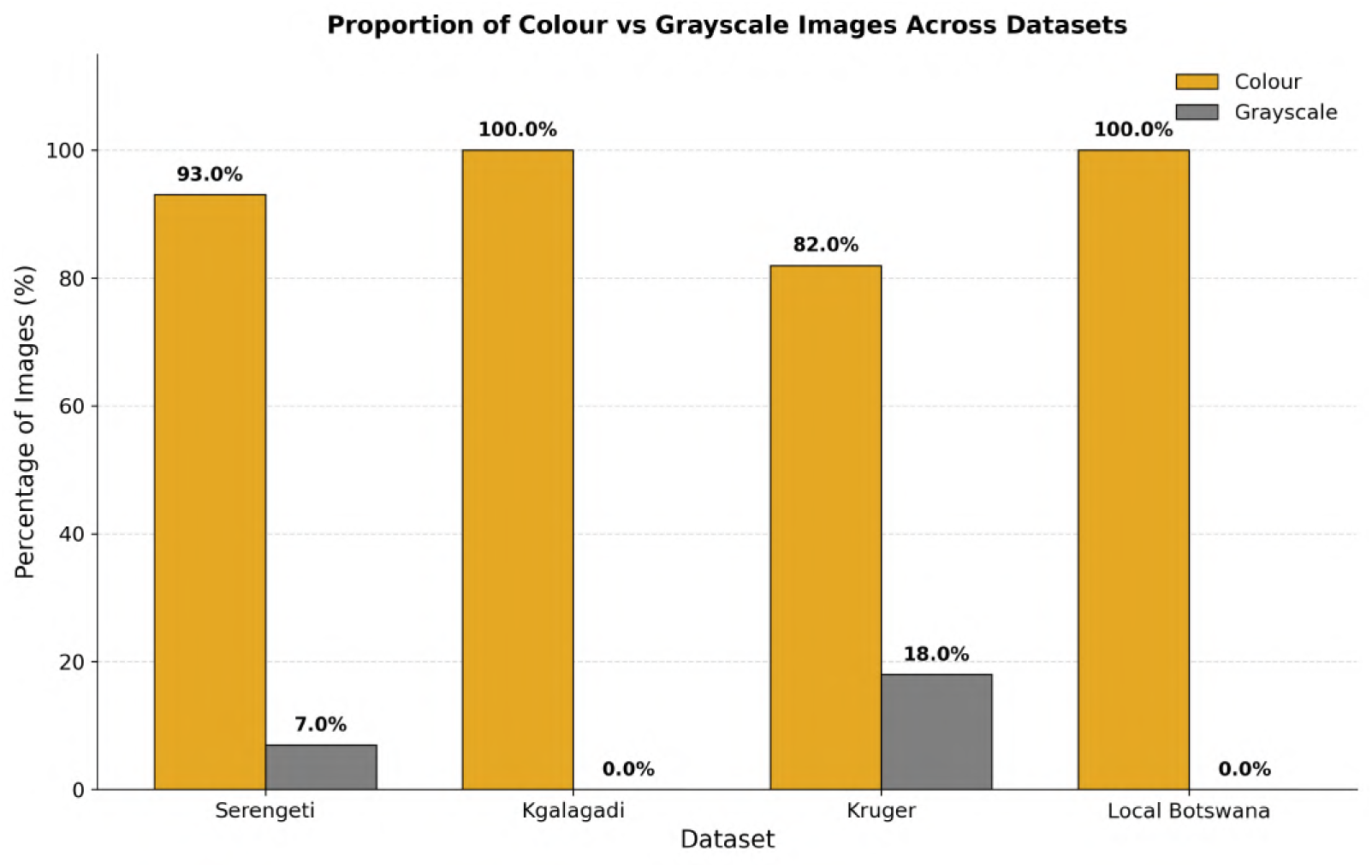
Proportion of colour versus grayscale images across all four datasets based on pixel-level channel analysis of 100 randomly sampled images per dataset. Kgalagadi and Local Botswana contain exclusively colour images. Kruger has the highest grayscale proportion at 18.0%. Percentages sum to 100% per dataset.

Day versus night capture proportions were assessed separately from colour versus grayscale analysis (Figure 6). For Serengeti, day versus night was determined from the timestamp column in the full Snapshot Serengeti Seasons 1–11 annotations metadata file, which contains 2,659,222 unique captures after deduplication by capture identifier. For Kgalagadi and Kruger, day versus night was determined from the `capture_time_local` column in the Snapshot Safari report CSV files. Daytime was defined as captures between 06:00 and 18:00 local time and nighttime as captures outside this window (Tabak et al., 2019). Serengeti has 82.7% daytime and 17.3% nighttime captures. Kgalagadi has 91.3% daytime and 8.7% nighttime captures across 3,626 sequences. Kruger has the highest proportion of nighttime activity at 35.3%, with only 64.7% daytime captures across 4,758 sequences, consistent with the high diversity of nocturnal predators and prey in the mixed woodland savanna ecosystem. All local Botswana photographs were collected during daylight hours and contain no nighttime captures.

**Figure 6.**
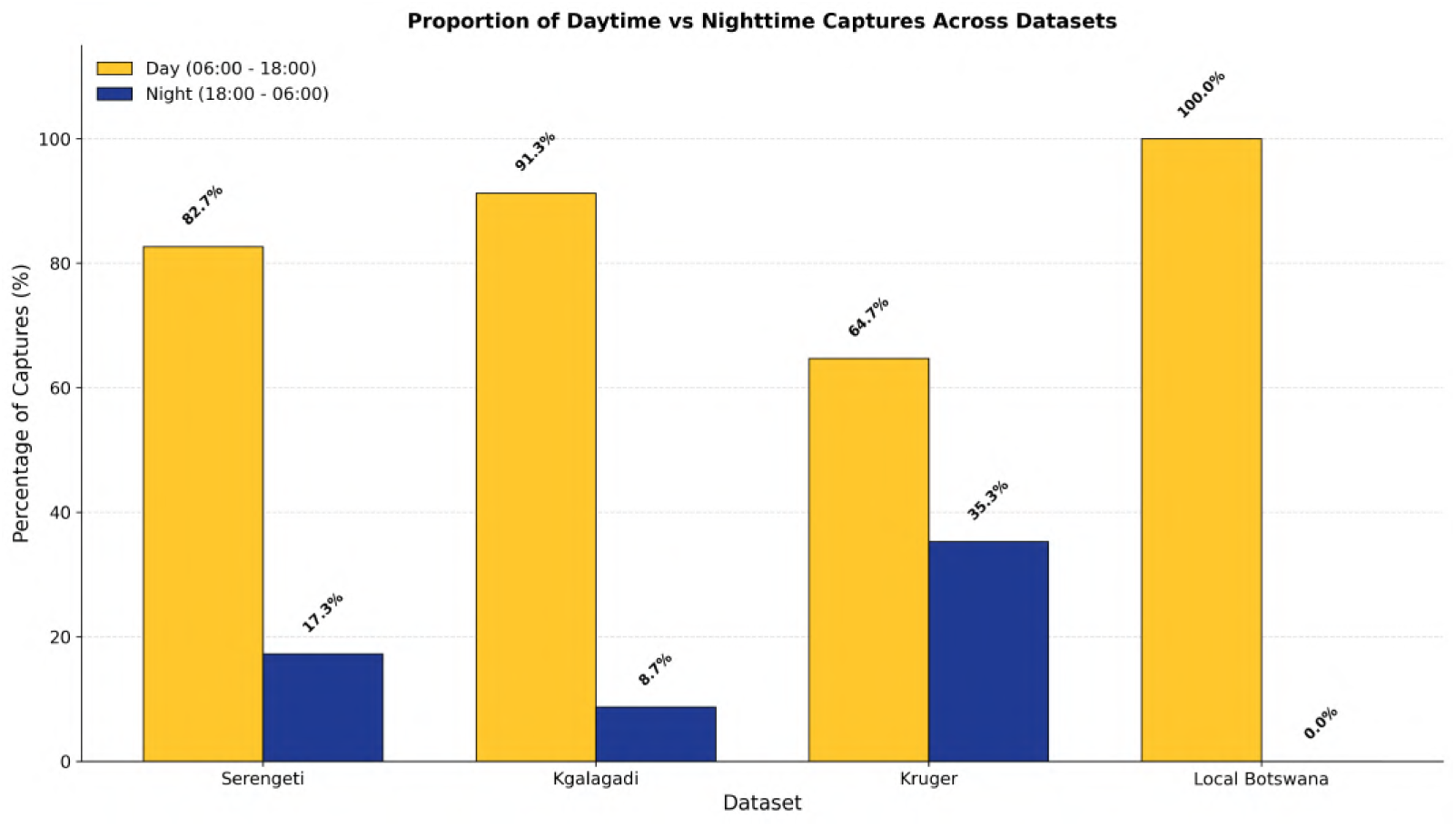
Proportion of daytime versus nighttime captures across all four datasets determined from actual timestamp metadata. For Serengeti, timestamps were extracted from the full Snapshot Serengeti Seasons 1–11 annotations file (*n* = 2,659,222 unique captures). For Kgalagadi and Kruger, timestamps were extracted from the Snapshot Safari report CSV files. Kruger has the highest nighttime proportion at 35.3%, substantially higher than its 18.0% grayscale proportion from pixel analysis, confirming that pixel-level grayscale analysis underestimates true nighttime activity rates. Local Botswana images are confirmed 100% daytime.

The difference between colour versus grayscale proportions and day versus night proportions is particularly evident for Serengeti, where pixel-level analysis estimated 7.0% nighttime captures but real timestamp data reveals 17.3%, and for Kruger where pixel analysis estimated 18.0% grayscale but real timestamps show 35.3% nighttime captures. This discrepancy confirms that colour versus grayscale pixel analysis substantially underestimates true nighttime capture rates when cameras retain partial colour sensitivity in low light conditions (Vyskočil and Picek, 2025). This distinction is directly relevant to Experiment 7, which evaluates model robustness under grayscale conditions. The higher true nighttime proportions suggest that real-world deployment performance would be lower than the colour-only experiment results indicate, particularly for Kruger.

Class distribution is well balanced in the Serengeti training set, where most species have approximately 3,000 images with the exception of hare and waterbuck which are capped at 1,000 images each and jackal at 2,500 (Figure 7). The cross-domain test sets are considerably more imbalanced. In Kruger, impala dominates with 585 images representing approximately 40% of all test images, while jackal has only 9 images (Figure 9). In Kgalagadi, ostrich is the most frequent species with 46 images while eland has only 14 and wildebeest only 13 (Figure 8). In the local Botswana dataset, elephant dominates with 177 images representing 56.0% of all photographs, reflecting the research team’s primary field sites near waterholes in northern Botswana (Figure 10). This severe imbalance across all test sets reinforces the choice of macro F1 as the primary evaluation metric in this study, as overall accuracy would be disproportionately influenced by dominant species, masking poor performance on rare but ecologically important taxa (Norouzzadeh et al., 2018; Willi et al., 2019).

**Figure 7.**
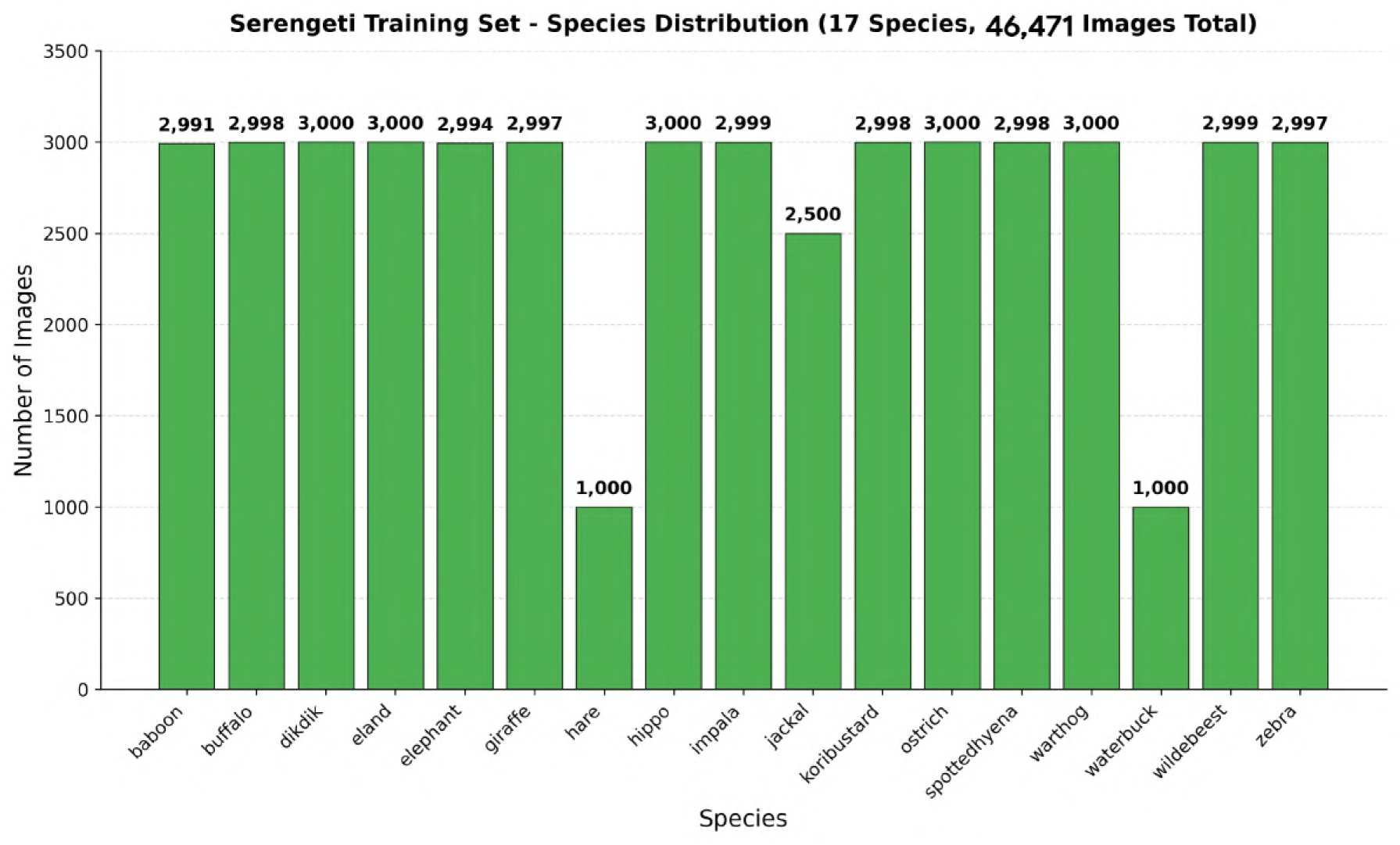
Species distribution in the Snapshot Serengeti dataset across 17 speciesto-talling 46,471 images (the full pre-split dataset). Most species are represented by approximately 3,000 images with the exception of hare and waterbuck at 1,000 each and jackal at 2,500, reflecting natural rarity constraints in the source dataset.

**Figure 8.**
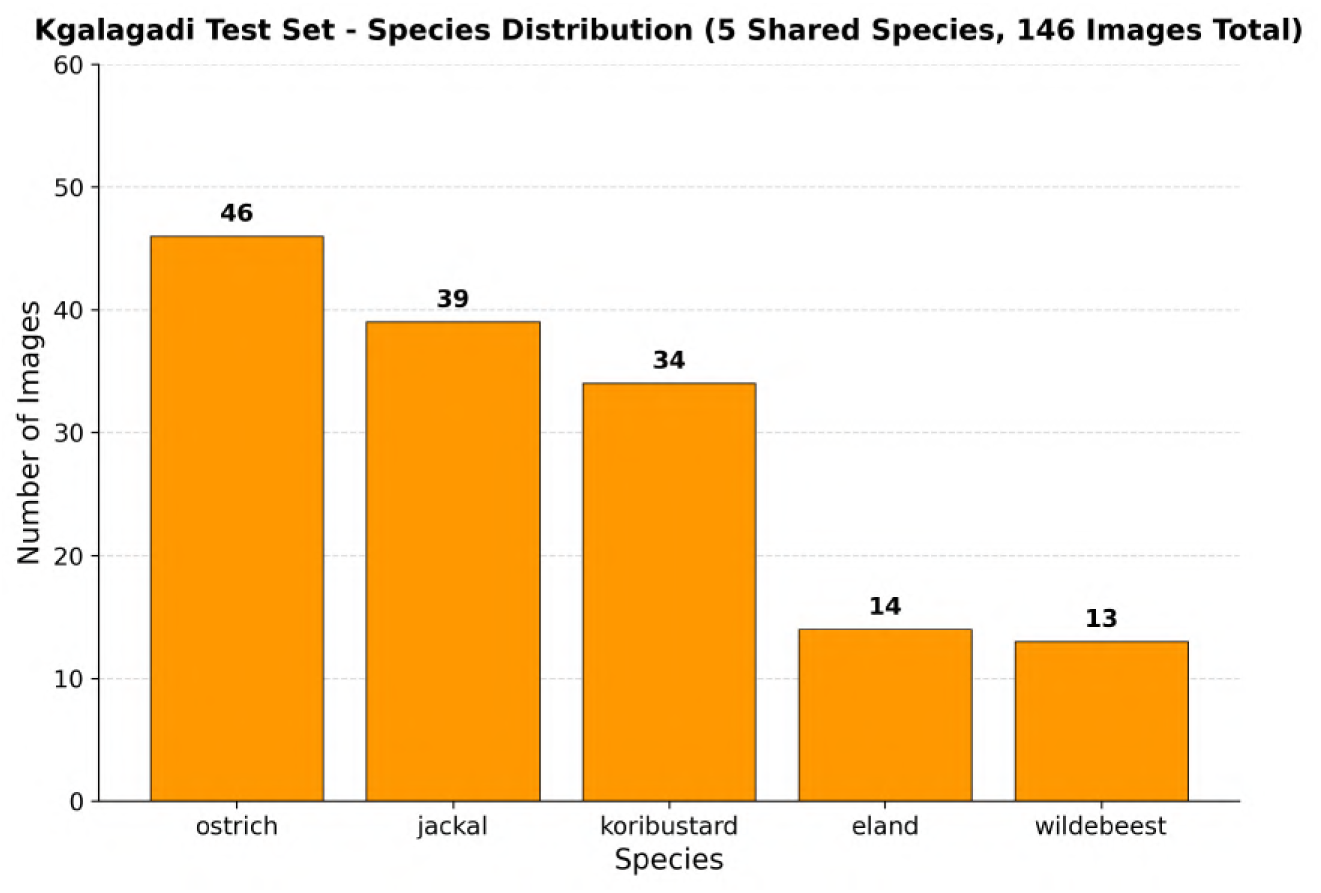
Species distribution in the Snapshot Kgalagadi test set across the 5 shared species retained after applying the minimum threshold of 9 images per species, totalling 146 test images. The dataset is moderately imbalanced with ostrich comprising 31.5% of all test images.

**Figure 9.**
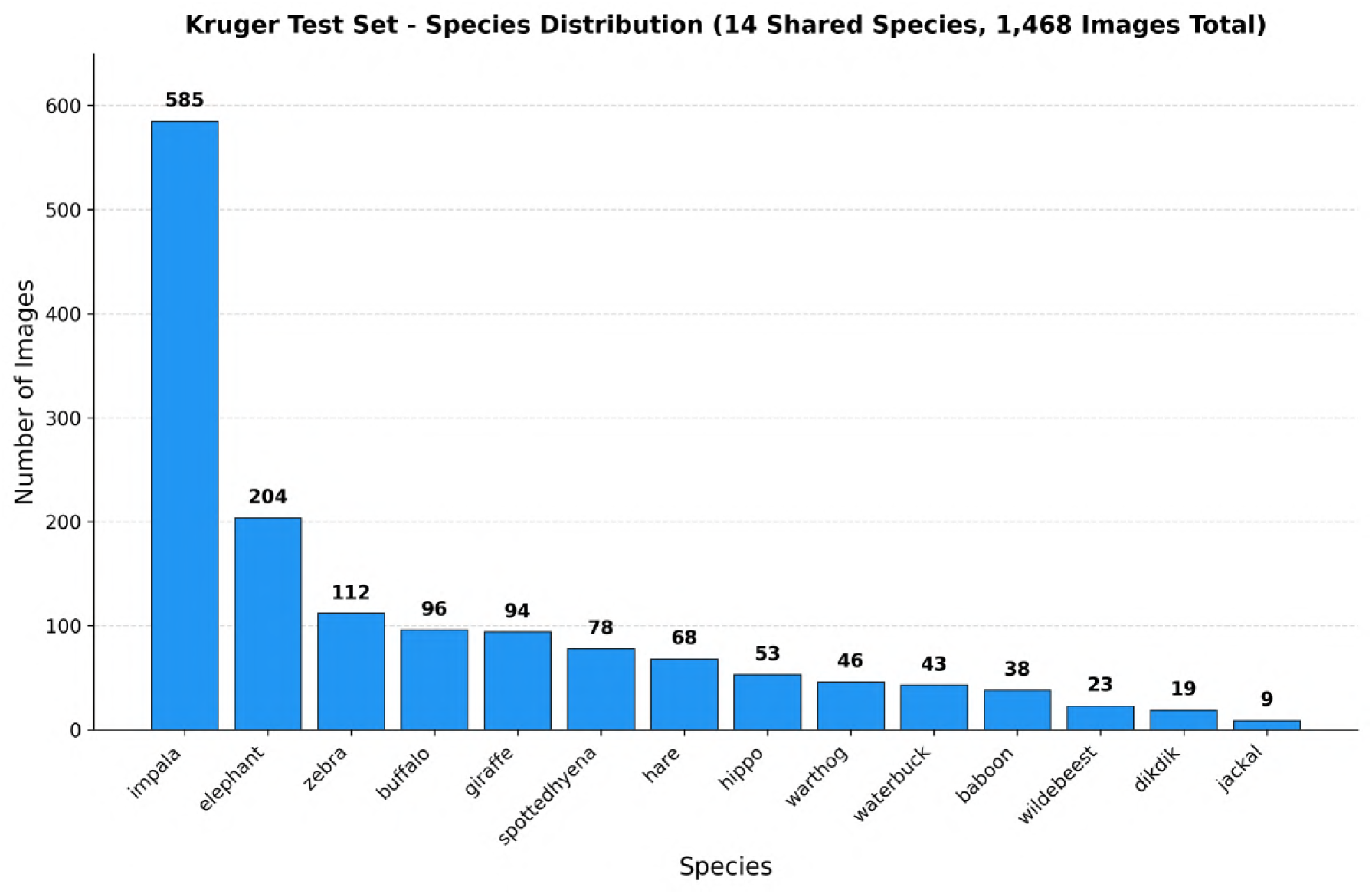
Species distribution in the Snapshot Kruger test set across the 14 shared species retained after filtering, totalling 1,468 test images. The dataset is heavily imbalanced with impala comprising approximately 40% of all test images and jackal represented by only 9 images.

**Figure 10.**
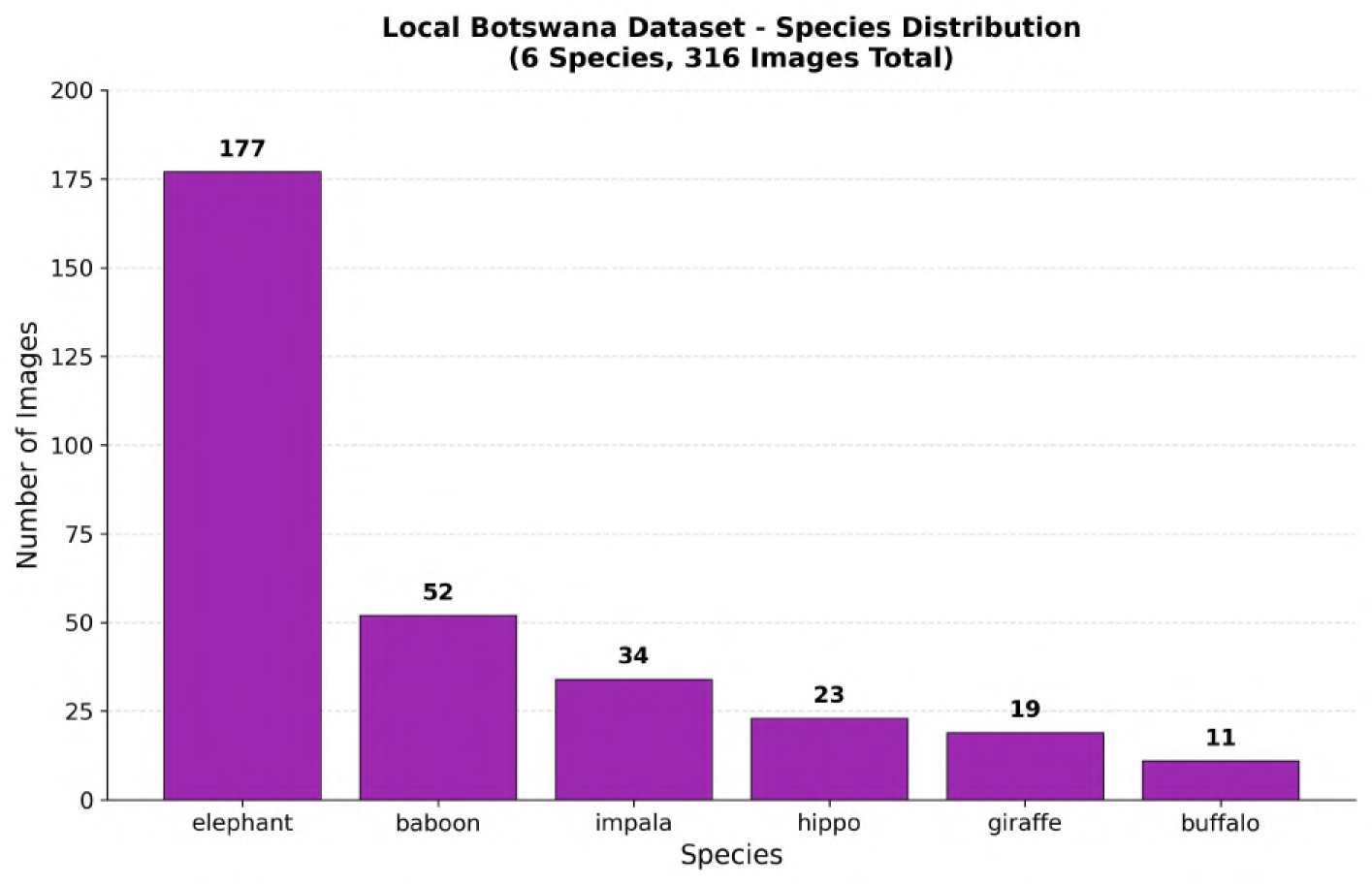
Species distribution in the Local Botswana wildlife photography dataset across 6 species totalling 316 images. Elephant dominates with 177 images (56.0%) re-flecting the research team’s primary field sites near waterholes in northern Botswana.

Mean image brightness differs across all four datasets (Figure 11). Kruger is the brightest at a mean pixel value of 122.8, followed by Kgalagadi, while Serengeti is the darkest at 110.6. Local Botswana images have the tightest brightness distribution, reflecting consistent daytime photography conditions with a single photographer. Serengeti exhibits the greatest brightness variability, reflecting the diversity of lighting conditions across its eleven seasons of continuous operation. These brightness differences represent a measurable visual domain shift between the training and test sets that is independent of species composition and geographic location (Schneider et al., 2020).

**Figure 11.**
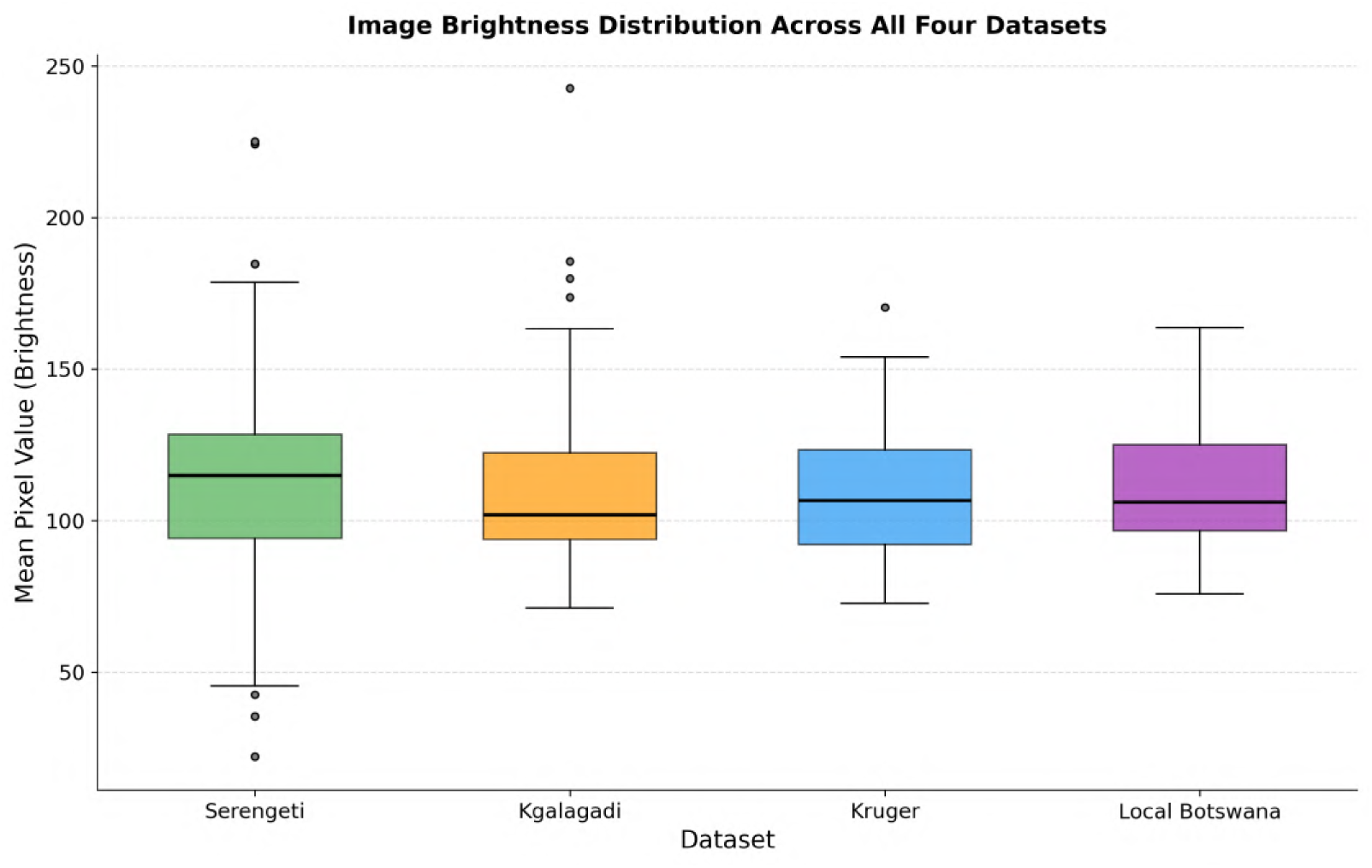
Distribution of mean pixel brightness values across 100 randomly sampled images per dataset for all four datasets. Local Botswana images have the tightest brightness distribution reflecting consistent daytime photography conditions. Serengeti exhibits the greatest brightness variability reflecting diverse lighting conditions across eleven seasons of continuous operation (Schneider et al., 2020).

### Models

We evaluate three model architectures that represent fundamentally different approaches to wildlife species classification. The first is a supervised fine-tuned transformer that requires labelled training data from the source domain. The second is a retrieval-based zero-shot approach that requires no training at all. The third is a true zero-shot foundation model pretrained specifically on biological imagery.

#### BEiTV2 — Supervised Fine-Tuned Baseline

BEiTV2 is a vision transformer model introduced by Peng et al. (2022) that uses masked image modelling with vector-quantised visual tokenizers as its pretraining objective. Rather than predicting raw pixel values from masked patches as earlier masked autoencoders do, BEiTV2 trains the model to predict discrete visual tokens derived from a learned visual tokenizer, encouraging the model to develop high-level semantic representations rather than low-level texture features. This approach has been shown to produce stronger and more generalisable visual features compared to earlier self-supervised vision transformers. Vyskočil and Picek (2025) demonstrated in camera trap experiments that BEiTV2 outperforms standard CNN architectures, particularly for rare tail species, and shows less location-specific overfitting than convolutional models. In our study, BEiTV2 serves as the supervised baseline. We fine-tune the pretrained BEiTV2 model on the Snapshot Serengeti training data by replacing the classification head with a new linear layer matching the number of training species and optimising all parameters end-to-end. Pretrained weights are loaded from the `microsoft/beit-base-patch16-224-pt22k` checkpoint on HuggingFace, which provides BEiTV2 weights pretrained on ImageNet-22k and serves as the starting point for downstream fine-tuning on the Serengeti training data. The full fine-tuning pipeline is illustrated in Figure 12.

**Figure 12.**
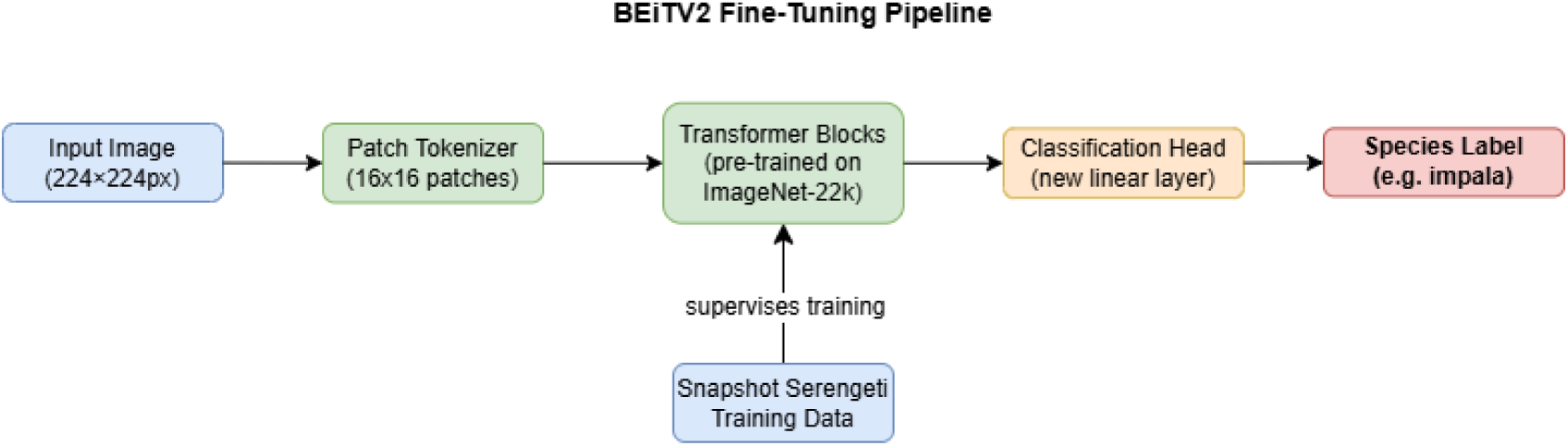
BEiTV2 fine-tuning pipeline. The input image is divided into 16*×*16 patches and passed through the pretrained transformer blocks, whose weights are updated end-to-end during fine-tuning on the Snapshot Serengeti training split (32,524 images, 17 species). The original classification head is replaced with a new linear layer matching the number of training species and trained from scratch.

#### DINOv2 with FAISS — Retrieval-Based Zero-Shot

DINOv2 is a self-supervised vision transformer introduced by Oquab et al. (2023) that produces high-quality all-purpose visual features without using any labels or annotations during pretraining. The model is trained on a curated dataset of 142 million images and produces feature representations that generalise strongly across diverse image domains without requiring any fine-tuning. In this study, we use DINOv2 as a feature extractor in a retrieval-based classification pipeline. We extract DINOv2 embeddings for all images in the Serengeti training set and store them in a FAISS vector index, which allows efficient similarity search across large embedding databases (Douze et al., 2024). At inference time, the embedding of a test image is compared against the indexed training embeddings using cosine similarity, and the species label of the nearest neighbour in the embedding space is assigned as the predicted class. This approach requires no model training whatsoever and makes classification decisions purely based on visual feature similarity. We use the DINOv2 ViT-Large model with a patch size of 14, which produces feature vectors of dimension 1024. The retrieval pipeline is illustrated in Figure 13.

**Figure 13.**
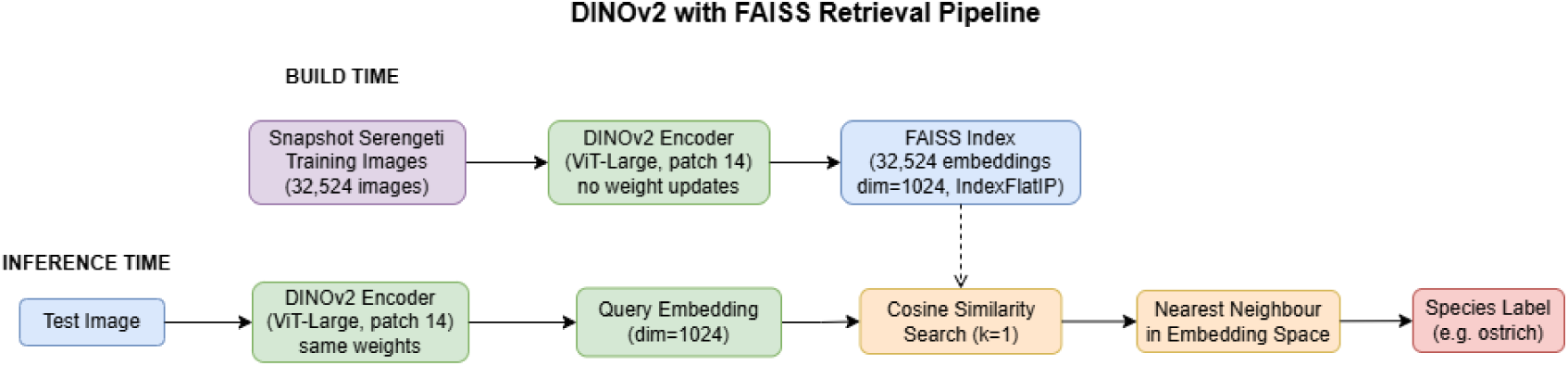
DINOv2 with FAISS retrieval pipeline. At build time, DINOv2 (ViT-Large, patch 14) extracts 1024-dimensional embeddings from all 32,524 Snapshot Serengeti training-split images and stores them in a FAISS index using the `IndexFlatIP` structure. At inference time, the same encoder generates a query embedding for the test image, which is compared against all stored embeddings via cosine similarity. The species label of the single nearest neighbour (*k* = 1) in the embedding space is assigned as the prediction. No model weights are updated at any point.

#### BioCLIP — True Zero-Shot

BioCLIP is a vision-language foundation model introduced by Stevens et al. (2024) that was pretrained specifically on biological imagery using a contrastive learning objective similar to CLIP but applied to the tree of life taxonomy. The model was trained on the TreeOfLife-10M dataset, a large collection of biological images paired with structured taxonomic descriptions, enabling it to recognise species across the full range of biological diversity including rare and underrepresented taxa. BioCLIP is used in this study as a true zero-shot model, meaning it receives no training data from any of our datasets and classifies species purely by matching image features against text embeddings of species names using the standard CLIP zero-shot inference protocol. Specifically, for each species in the target dataset, we generate a text prompt of the form “a photo of a [species name]” and compute text embeddings. This prompt template follows the standard zero-shot inference protocol established by Radford et al. (2021), who demonstrated that embedding the class name within a natural language sentence consistently outperforms using bare class labels alone, as the sentence context better aligns the text embedding with the visual representations learned during contrastive pretraining. At inference time, the image embedding is matched against all species text embeddings and the species with the highest cosine similarity is selected as the prediction. Pretrained weights are available at `imageomics/bioclip` on HuggingFace. The zero-shot inference pipeline is illustrated in Figure 14.

**Figure 14.**
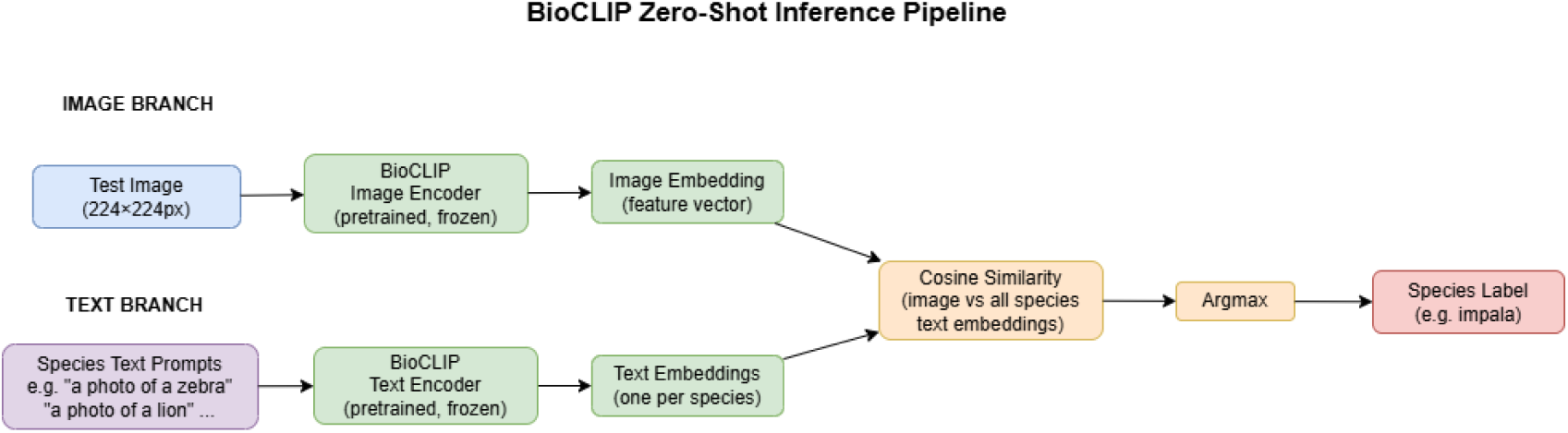
BioCLIP zero-shot inference pipeline. Two parallel branches produce embeddings that are compared via cosine similarity. In the image branch, the test image is encoded by the BioCLIP image encoder into a feature vector. In the text branch, species name prompts of the form “a photo of a [species name]” are encoded by the BioCLIP text encoder into one text embedding per species. The species whose text embedding has the highest cosine similarity to the image embedding is selected as the prediction via argmax. No training data is used at any point — BioCLIP operates in a purely zero-shot manner.

### Experiments

We design eight experiments to systematically investigate geographic domain shift across our datasets and to evaluate the effect of various preprocessing and data decisions on model performance.

#### Experiment 1 — In-Domain Baseline

In this experiment, all three models are evaluated on Snapshot Serengeti data only using a standard train/validation/test split. BEiTV2 is fine-tuned on the Serengeti training split. DINOv2 uses the Serengeti training images as the retrieval index without any weight updates. BioCLIP classifies Serengeti test images in a purely zero-shot manner using text prompts, with no exposure to any training images. This provides the upper-bound performance reference for BEiTV2 and establishes the performance level achievable when there is no domain shift. The results from this experiment serve as the comparison point for all subsequent cross-dataset experiments.

#### Experiment 2 — Serengeti to Kgalagadi

Models are evaluated directly on Snapshot Kgalagadi without any additional adaptation. This experiment measures the performance drop caused by the geographic shift from East African savanna to the arid Southern African Kalahari ecosystem. For BioCLIP and DINOv2 with FAISS, which require no training, the Serengeti data still serves as the retrieval database or the source of zero-shot species name prompts.

#### Experiment 3 — Serengeti to Kruger

The same inference procedure as Experiment 2 is applied, this time evaluating all three models on Snapshot Kruger. While Experiment 2 measures the performance drop caused by transferring from East African savanna to the arid Kalahari desert ecosystem, Experiment 3 measures the drop when transferring to the mixed woodland savanna of Kruger National Park. Comparing the two experiments allows us to assess whether the severity of geographic domain shift is proportional to the ecological distance between source and target ecosystems, with Kruger expected to represent a less extreme shift than Kgalagadi given its greater species overlap with Serengeti.

#### Experiment 4 — Serengeti to Local Botswana Images

Models are evaluated on the locally collected Botswana photography dataset. This experiment represents the most challenging domain shift scenario because the target images were not collected using camera traps and differ substantially in image format, viewing angle, and capture conditions from the training data.

#### Experiment 5 — Data Scaling Analysis

To investigate how much training data is needed to achieve acceptable performance, we train BEiTV2 on subsets of the Serengeti training split representing 1%, 5%, 25%, 50%, and 100% of the available training images. Fractions were chosen following Bevan et al. (2026) to span the extreme degradation regime (1%, 5%), the transition region (25%), and the saturation regime (50%, 100%) documented in their analysis of camera trap data scaling. Performance is measured on the in-domain Serengeti test set and on all three cross-domain test sets (Kgalagadi, Kruger, and Local Botswana) at each data level. This experiment provides practical guidance for conservation practitioners who may not have access to the full Serengeti dataset and need to understand the minimum data requirement for useful model performance (Bevan et al., 2026; Schneider et al., 2020; Whytock et al., 2021).

#### Experiment 6 — MegaDetector Preprocessing

MegaDetector is a widely used object detector developed by Beery et al. (2019b) that locates animals within camera trap images and produces bounding box crops removing background clutter. We run all three models on both the full raw images and on MegaDetector-cropped versions of the same images, and compare performance across both conditions. This allows us to evaluate whether removing background information before classification improves performance and reduces the location-specific overfitting that Beery et al. (2018) identified as a primary cause of domain shift.

#### Experiment 7 — Grayscale versus Colour Images

Camera traps frequently capture images in black and white, particularly at night using infrared sensors. To assess whether models are relying on colour information that may not always be available in the field, we convert all images to grayscale and repeat the classification experiments. Comparing accuracy between colour and grayscale conditions reveals the extent to which colour is a useful feature for each model and informs deployment decisions in settings where night-time or infrared images are common.

#### Experiment 8 — Per-Species Transfer Analysis

For each cross-dataset experiment, we compute per-species accuracy, precision, recall, and F1 score to identify which individual species suffer the most from geographic domain shift. Species are then grouped by characteristics such as whether they appear in both the source and target dataset, how many training images they have in Serengeti, and whether they are visually similar to other species. This analysis provides ecologically meaningful insights about which taxa are hardest to classify across regions and informs which species may need additional locally collected data for reliable monitoring.

### Evaluation Metrics

We evaluate all models using five complementary metrics that capture different aspects of classification performance (Sokolova and Lapalme, 2009). Let *N* denote the total number of test images, *C* the number of species classes present in the test set, and *TP_c_*, *FP_c_*, *TN_c_*, and *FN_c_* denote the true positives, false positives, true negatives, and false negatives for class *c*, respectively.

Overall accuracy reports the proportion of correctly classified images across all species:

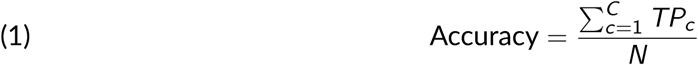

While accuracy is reported for direct comparison with prior work, it can be misleading when class distributions are highly imbalanced, as common species dominate the aggregate score (Norouzzadeh et al., 2018; Sokolova and Lapalme, 2009).

Precision for class *c* measures the proportion of predicted positives that are correct (Sokolova and Lapalme, 2009):

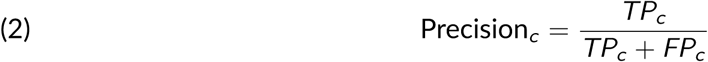

Recall for class *c* measures the proportion of true positives that are correctly identified (Sokolova and Lapalme, 2009):

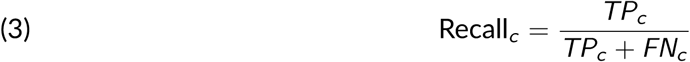

F1 score for class *c* is the harmonic mean of precision and recall, providing a single balanced measure of a model’s performance on that class (Sokolova and Lapalme, 2009):

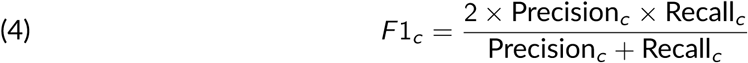

Macro F1 score computes the F1 score for each species independently and averages across all *C* classes present in the test set with equal weight regardless of class frequency (Sokolova and Lapalme, 2009):

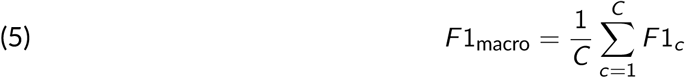

Macro F1 is adopted as the primary evaluation metric in this study because it treats all species equally regardless of their frequency in the test set, which is critical for rare species that are ecologically significant but contribute little to overall accuracy. This directly addresses the concern raised by Norouzzadeh et al. (2018) regarding inflated accuracy scores on imbalanced camera trap datasets, where high overall accuracy can mask poor performance on rare but conservation-relevant species.

Wall-clock training time, inference time per image, and peak GPU memory are recorded for each model alongside the classification metrics. These are reported because deployment feasibility for conservation practitioners working in field settings depends on computational efficiency as well as predictive accuracy, and we want the experimental results to communicate both. All measurements are taken on the same hardware (NVIDIA GeForce RTX 3070 Laptop GPU with 8 GB VRAM, 16 GB system RAM).

#### Statistical significance testing

To assess whether observed differences in performance between models are statistically significant rather than due to chance, we apply McNemar’s test (McNemar, 1947) to pairwise comparisons of model predictions on each test set with Edwards’ continuity correction. McNemar’s test is appropriate for comparing the error rates of two classifiers evaluated on the same test set and operates on the contingency table of cases where one model predicts correctly and the other does not. Given classifiers *A* and *B*, the test statistic with continuity correction is:

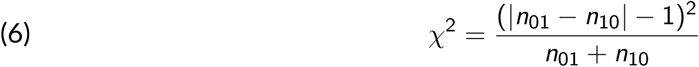

where *n*_01_ is the number of test images correctly classified by model *B* but not by model *A*, and *n*_10_ is the number correctly classified by model *A* but not by model *B*. This statistic follows a chi-squared distribution with one degree of freedom under the null hypothesis that both models have the same error rate (McNemar, 1947). A significance threshold of *α* = 0.05 is used throughout. The discordant odds ratio *n*_01_*/n*_10_ is reported alongside *p*-values as an effect size, with values greater than 1 indicating that model *B* wins more disagreements than model *A*. Wilson 95% confidence intervals (Wilson, 1927) are reported for all accuracy point estimates, following the convention used by Willi et al. (2019) and Tabak et al. (2020) in comparable wildlife camera-trap classification studies.

Domain Distance Scoring. To quantify the statistical distance between the source and target feature distributions, we compute the Maximum Mean Discrepancy (MMD) (Gretton et al., 2012) between the DINOv2 embedding spaces of each dataset pair. MMD is a nonparametric kernel-based metric that measures the distance between two probability distributions by comparing their mean embeddings in a reproducing kernel Hilbert space (RKHS). Given source distribution *P* and target distribution *Q*, the squared MMD with kernel *k* is defined as:

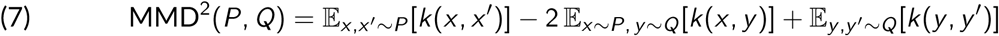

A value of zero indicates that the two distributions are identical, while larger values indicate greater distributional divergence (Gretton et al., 2012). MMD was selected over alternative domain distance metrics such as the Fréchet Inception Distance (FID) for two reasons. First, MMD operates directly on the DINOv2 feature embeddings already extracted during the retrieval pipeline, requiring no additional computational overhead. Second, MMD is a well-established, theoretically grounded metric with a closed-form estimator that is suitable for the sample sizes available in our test sets (Gretton et al., 2012). MMD is computed between Serengeti and Kgalagadi, Serengeti and Kruger, and Serengeti and Local Botswana, using a Radial Basis Function (RBF) kernel with bandwidth set to the median pairwise distance of the combined sample. The resulting MMD scores are reported alongside classification results in Section 4 to provide quantitative context for the magnitude of domain shift observed in each cross-regional experiment.

### Hyperparameters

Hyperparameters for BEiTV2 fine-tuning were initialized by adapting the task-adaptation framework of Peng et al. (2022) and modifying specific configuration thresholds to account for our smaller-scale camera-trap data and severe class imbalances, validated against the downstream optimization regimes explored by Vyskočil and Picek (2025). While Peng et al. (2022) utilize a base learning rate of 5 *×* 10^−4^ for full-scale ImageNet fine-tuning, we adopted a more conservative learning rate of 1 *×* 10^−4^ to ensure stable gradient updates and prevent representations from shifting too aggressively away from pretrained weights given our target ecosystem contexts.

A layer-wise learning rate decay of 0.65 was adopted directly from the base-sized ViT fine-tuning protocol of Peng et al. (2022), which applies progressively smaller learning rates to earlier transformer layers to preserve foundational feature representations while allowing the classification head and higher layers to fully adapt. Rather than using the large-batch regime of 1024 profiles detailed by Peng et al. (2022), we established an effective batch size of 64 (a micro-batch size of 16 accumulated over 4 steps) to accommodate local computational hardware limitations while ensuring sufficient variance smoothing.

To explicitly safeguard against background overfitting on constrained camera-trap sequences, we incorporated a dropout rate of 0.1 as a manual regularization constraint, contrasting with the bare stochastic depth configurations utilized in the original Peng et al. (2022) experiments. Weight decay was maintained at 0.05, and label smoothing was kept at 0.1 to systematically soften hard targets and soften overconfidence on minority-class labels. Our training duration was bounded at a maximum of 50 epochs with a 5-epoch linear warmup—shortened from the standard 100-epoch base schedule of Peng et al. (2022)—and tied to an early stopping criterion with a patience of 10 epochs based on validation performance. No systematic hyperparameter optimization sweep was conducted; all baseline variables were held static following initialization.

The finalized hyperparameters for fine-tuning BEiTV2 on the Snapshot Serengeti dataset are summarized in Table 2.

**Table 2.** BEiTV2 fine-tuning hyperparameters.

| Hyperparameter | Value | Short definition |
| --- | --- | --- |
| Optimiser | AdamW | The optimisation algorithm used to update model weights during training. |
| Base learning rate | $1 \times 10^{-4}$ | The initial step size controlling how much model weights change during each update. |
| Layer-wise LR decay | 0.65 | A strategy that applies smaller learning rates to lower layers and larger to higher layers during fine-tuning. |
| Batch size | 16 | The number of training images processed in one forward and backward pass before updating the model. |
| Gradient accumulation | 4 steps | The number of forward passes accumulated before one weight update, giving an effective batch size of 64. |
| Maximum epochs | 50 | The maximum number of complete passes through the training dataset. |
| Early stopping patience | 10 epochs | The number of epochs to wait for validation improvement before stopping training early. |
| Weight decay | 0.05 | A regularisation term that discourages overly large model weights to reduce overfitting. |
| Warmup epochs | 5 | The number of initial epochs during which the learning rate is gradually increased to its target value. |
| Input image resolution | 224×224 px | The size to which all input images are resized before being passed into the model. |
| Dropout rate | 0.1 | The proportion of neurons randomly deactivated during training to improve generalisation. |
| Label smoothing | 0.1 | A regularisation technique that softens hard class labels to reduce overconfidence in predictions. |
| Mixed precision | fp16 | Half-precision floating point training that reduces GPU memory usage without significant loss in accuracy. |

#### DINOv2 with FAISS Hyperparameters

DINOv2 requires no training so there are no optimisation hyperparameters. The relevant settings that control the behaviour of the retrieval pipeline are shown in Table 3.

**Table 3.** DINOv2 with FAISS retrieval pipeline settings.

| Hyperparameter | Value | Short definition |
| --- | --- | --- |
| DINOv2 model variant | ViT-Large, patch 14 | The specific pretrained DINOv2 backbone used to extract visual features from images. |
| Embedding dimension | 1024 | The size of the feature vector produced for each image. |
| FAISS index type | IndexFlatIP | The similarity search structure used to perform exact nearest-neighbour retrieval based on inner product. |
| Feature normalisation | L2 before indexing | A preprocessing step that scales feature vectors to unit length before storage and comparison. |
| Similarity metric | Cosine similarity | The measure used to determine how visually similar two image embeddings are. |
| Nearest neighbours ($k$) | 1 | The number of closest training images considered when assigning the predicted species label. |
| Input image resolution | 224×224 px | The size to which all input images are resized before feature extraction. DINOv2-Large supports this size via positional-embedding interpolation; this is below the 518×518 most commonly used in DINOv2-L benchmarks. |

The FAISS `IndexFlatIP` index performs exact nearest-neighbour search using inner product on L2-normalised vectors, which is mathematically equivalent to cosine similarity (Douze et al., 2024). The *k*= 1 setting means the predicted species label is taken directly from the single most similar training image in the embedding database, following the approach used by Vyskočil and Picek (2025) for camera trap retrieval classification.

#### BioCLIP Inference Hyperparameters

BioCLIP is used entirely in inference mode with no parameter updates. The settings that control its zero-shot classification behaviour are shown in Table 4.

**Table 4.**
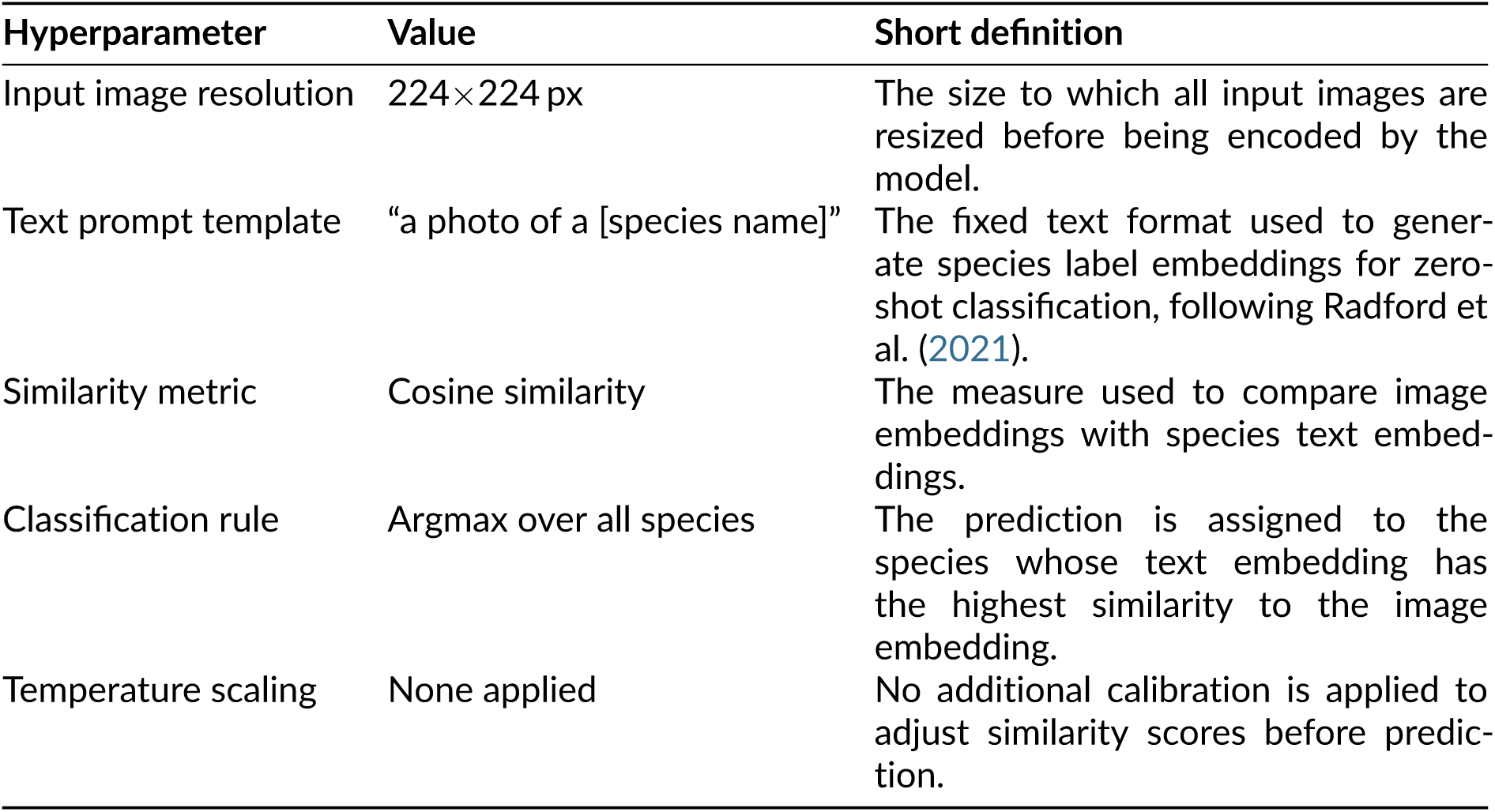
BioCLIP zero-shot inference settings.

| Hyperparameter | Value | Short definition |
| --- | --- | --- |
| Input image resolution | 224×224 px | The size to which all input images are resized before being encoded by the model. |
| Text prompt template | “a photo of a [species name]” | The fixed text format used to generate species label embeddings for zero-shot classification, following Radford et al. (2021). |
| Similarity metric | Cosine similarity | The measure used to compare image embeddings with species text embeddings. |
| Classification rule | Argmax over all species | The prediction is assigned to the species whose text embedding has the highest similarity to the image embedding. |
| Temperature scaling | None applied | No additional calibration is applied to adjust similarity scores before prediction. |

The prompt template “a photo of a [species name]” follows the standard zero-shot inference protocol established by Radford et al. (2021) for CLIP-style models, where the class name is embedded in a natural language sentence to produce a richer text representation than using the bare class label alone. The species with the highest cosine similarity between its text embedding and the query image embedding is selected as the predicted label.

## Results

### Experiment 1 — In-Domain Baseline (Serengeti)

Table 5 presents the performance of all three models evaluated on the in-domain Snapshot Serengeti test set (*n* = 6,982 images, 17 species). This experiment establishes the upper-bound performance reference for each model under conditions where no geographic domain shift is present, providing the baseline against which all cross-regional results are compared. Overall accuracy is reported with Wilson 95% confidence intervals, and macro F1 serves as the primary metric as it weights all species equally regardless of class frequency.

**Table 5.** Experiment 1: in-domain performance on the Snapshot Serengeti test set (*n* = 6,982 images, 17 species). Accuracy is reported with Wilson 95% confidence intervals. Inference time is measured in milliseconds per image on an NVIDIA GeForce RTX 3070.

| Model | Accuracy % (95% CI) | Macro F1 (%) | Inference (ms/img) |
| --- | --- | --- | --- |
| DINOv2 + FAISS | 90.52 [89.81, 91.18] | 90.65 | 0.48 |
| BEiT<sub>V2</sub> | 83.62 [82.73, 84.46] | 83.53 | 9.47 |
| BioCLIP | 46.76 [45.59, 47.93] | 47.53 | 9.96 |

DINOv2 with FAISS achieved the strongest in-domain performance with a top-1 accuracy of 90.52% (95% CI: 89.81–91.18%) and a macro F1 score of 90.65%, despite requiring no model training whatsoever and classifying images purely through nearest-neighbour retrieval in the embedding space. The close agreement between accuracy and macro F1 (90.52% and 90.65%) indicates performance is consistent across both common and rare species. The inference time of 0.48 ms per image reflects the efficiency of the FAISS index search rather than a full neural network forward pass.

BEiTV2, the supervised fine-tuned baseline, achieved 83.62% accuracy (95% CI: 82.73–84.46%) and 83.53% macro F1 after training on the Serengeti training split, with an inference time of 9.47 ms per image. The close alignment between accuracy and macro F1 again indicates balanced performance across species.

BioCLIP, the true zero-shot model that received no Serengeti data at any point, achieved 46.76% accuracy (95% CI: 45.59–47.93%) and 47.53% macro F1. This result is well above the random chance baseline of 5.88% for a 17-class classification task. The gap between DINOv2 and BioCLIP under in-domain conditions reflects the difference between the two zero-shot approaches: DINOv2 has access to 32,524 Serengeti training-split images as a retrieval index at inference time, whereas BioCLIP relies entirely on its biological pretraining and text prompts with no access to any Serengeti imagery.

Pairwise McNemar’s tests with Edwards’ continuity correction confirm that all three observed performance differences are statistically significant at *α* = 0.05 (Table 6).

**Table 6.** Experiment 1: pairwise McNemar’s test results on the Serengeti in-domain test set. The discordant odds ratio (OR) is defined as *n*_01_*/n*_10_, where *n*_01_ is the number of test images correctly classified by model B but not by A, and *n*_10_ the reverse; values greater than 1 indicate that B wins more disagreements than A.

| Model A | Model B | $n_{01}$ | $n_{10}$ | $\chi^2$ | $p$-value | OR |
| --- | --- | --- | --- | --- | --- | --- |
| BEiT2 | DINOv2 + FAISS | 739 | 257 | 232.29 | $< 0.001$ | 2.88 |
| BEiT2 | BioCLIP | 266 | 2839 | 2130.49 | $< 0.001$ | 0.09 |
| DINOv2 + FAISS | BioCLIP | 94 | 3149 | 2876.01 | $< 0.001$ | 0.03 |

All three pairwise tests yielded *p <* 0.001. DINOv2 with FAISS significantly outperformed BEiTV2 (*χ*^2^ = 232.29): of the 996 test images on which the two models disagreed, DINOv2 was correct on 739 and BEiTV2 on 257, an odds ratio of 2.88 in DINOv2’s favour. BEiTV2 significantly outperformed BioCLIP (*χ*^2^ = 2,130.49), winning 2,839 of the 3,105 discordant cases against BioCLIP’s 266, which corresponds to BEiTV2 being correct approximately 10.7 times as often as BioCLIP on disagreements. DINOv2 with FAISS displayed the largest performance gap over BioCLIP (*χ*^2^ = 2,876.01), winning 3,149 of 3,243 discordant cases, equivalent to DINOv2 being correct approximately 33.5 times as often.

### Experiment 2 — Serengeti to Kgalagadi

Table 7 presents the performance of all three models on Snapshot Kgalagadi (*n* = 146 images, 5 species) without any adaptation to the target domain. The Maximum Mean Discrepancy between Serengeti and Kgalagadi, computed using DINOv2 ViT-Large CLS token embeddings with an RBF kernel and median heuristic bandwidth (*σ* = 1.2893), was MMD^2^ = 0.0706, computed over *n* = 1,000 Serengeti and *m* = 146 Kgalagadi images.

**Table 7.** Experiment 2: cross-domain performance on Snapshot Kgalagadi (*n* = 146 images, 5 species) after training on Snapshot Serengeti only. Accuracy is reported with Wilson 95% confidence intervals. Performance drop in percentage points (pp) is measured relative to Experiment 1 accuracy.

| Model | Accuracy % (95% CI) | Macro F1 (%) | Acc. Drop (pp) |
| --- | --- | --- | --- |
| DINOv2 + FAISS | 77.40 [69.96, 83.43] | 79.81 | −13.12 |
| BioCLIP | 61.64 [53.55, 69.14] | 59.62 | +14.88 |
| BEiT2 | 28.08 [21.43, 35.86] | 29.30 | −55.54 |

DINOv2 with FAISS achieved 77.40% accuracy (95% CI: 69.96–83.43%) and 79.81% macro F1, the strongest of the three models on Kgalagadi and a drop of 13.12 percentage points in accuracy from the in-domain baseline. The macro F1 of 79.81% confirms that this performance is consistent across all five species rather than driven by any single dominant class.

BEiTV2 dropped from 83.62% in-domain to 28.08% (95% CI: 21.43–35.86%) on Kgalagadi, a fall of 55.54 percentage points in accuracy and 54.23 percentage points in macro F1. The near-identical drops in both metrics indicate that the degradation is consistent across all five species rather than concentrated on any single class.

BioCLIP showed an apparent accuracy increase of 14.88 percentage points on Kgalagadi compared to its in-domain Serengeti score, with a final accuracy of 61.64% (95% CI: 53.55–69.14%) and macro F1 of 59.62%. The Kgalagadi test set contains only 5 species compared to 17 in Serengeti, making the classification task substantially simpler in terms of the number of possible labels. The macro F1 score of 59.62% remains substantially below DINOv2’s 79.81% on the same test set.

Pairwise McNemar’s tests confirm that all three model differences on Kgalagadi are statistically significant at *α* = 0.05 (Table 8).

**Table 8.** Experiment 2: pairwise McNemar’s test results on Snapshot Kgalagadi (*n* = 146).

| Model A | Model B | $n_{01}$ | $n_{10}$ | $\chi^2$ | $p$-value | OR |
| --- | --- | --- | --- | --- | --- | --- |
| BEiT2 | DINOv2 + FAISS | 76 | 4 | 63.01 | $< 0.001$ | 19.00 |
| BEiT2 | BioCLIP | 62 | 13 | 30.72 | $< 0.001$ | 4.77 |
| DINOv2 + FAISS | BioCLIP | 10 | 33 | 11.26 | $< 0.001$ | 0.30 |

DINOv2 with FAISS significantly outperformed BEiTV2 (*χ*^2^ = 63.01, *p <* 0.001): of 80 discordant cases, DINOv2 was correct on 76 and BEiTV2 on 4, an odds ratio of 19.00 in DINOv2’s favour. BioCLIP also significantly outperformed BEiTV2 (*χ*^2^ = 30.72, *p <* 0.001): of 75 discordant cases, BioCLIP was correct on 62 and BEiTV2 on 13. DINOv2 with FAISS significantly outperformed BioCLIP on Kgalagadi (*χ*^2^ = 11.26, *p <* 0.001): of 43 discordant cases, DINOv2 was correct on 33 and BioCLIP on 10, equivalent to DINOv2 being correct approximately 3.3 times as often as BioCLIP on disagreements.

### Experiment 3 — Serengeti to Kruger

Table 9 presents the performance of all three models on Snapshot Kruger (*n* = 1,468 images, 14 species). Kruger shares 14 species with Serengeti compared to 5 for Kgalagadi. The Maximum Mean Discrepancy between Serengeti and Kruger, computed using DINOv2 ViT-Large CLS token embeddings with an RBF kernel and median heuristic bandwidth (*σ* = 1.2831), was MMD^2^ = 0.0348, computed over *n* = 1,000 Serengeti and *m* = 1,468 Kruger images. This is the smallest distributional distance of the three target datasets.

**Table 9.**
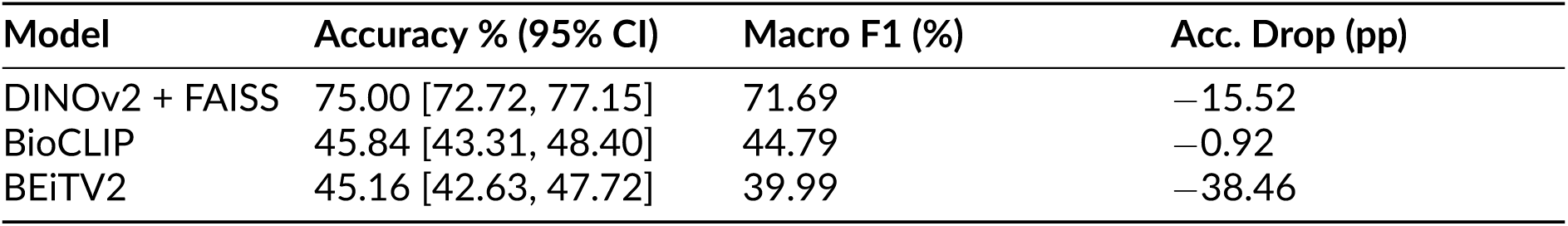
Experiment 3: cross-domain performance on Snapshot Kruger (*n* = 1,468 images, 14 species) after training on Snapshot Serengeti only. Accuracy is reported with Wilson 95% confidence intervals. Performance drop in percentage points (pp) is measured relative to Experiment 1 accuracy.

DINOv2 with FAISS again achieved the highest performance at 75.00% accuracy (95% CI: 72.72–77.15%) and 71.69% macro F1. The gap between accuracy (75.00%) and macro F1 (71.69%) reflects the heavier class imbalance of the Kruger test set where impala alone constitutes approximately 40% of test images. The drop from in-domain performance is 15.52 percentage points.

BEiTV2 dropped to 45.16% accuracy (95% CI: 42.63–47.72%) and 39.99% macro F1, a fall of 38.46 percentage points in accuracy from in-domain performance. The gap between accuracy (45.16%) and macro F1 (39.99%) indicates lower performance on less frequent species, with rare species showing variable per-class recall.

BioCLIP achieved 45.84% accuracy (95% CI: 43.31–48.40%) and 44.79% macro F1 on Kruger, a change of only 0.92 percentage points in accuracy from in-domain. The near-equal accuracy and macro F1 scores indicate consistent performance across all 14 Kruger species.

Pairwise McNemar’s tests reveal a mixed pattern of statistical significance on Kruger (Table 10).

**Table 10.** Experiment 3: pairwise McNemar’s test results on Snapshot Kruger.

| Model A | Model B | $n_{01}$ | $n_{10}$ | $\chi^2$ | $p$-value | OR |
| --- | --- | --- | --- | --- | --- | --- |
| BEiT2 | DINOv2 + FAISS | 467 | 29 | 385.02 | $< 0.001$ | 16.10 |
| BEiT2 | BioCLIP | 237 | 227 | 0.18 | 0.676 | 1.04 |
| DINOv2 + FAISS | BioCLIP | 37 | 465 | 363.21 | $< 0.001$ | 0.08 |

DINOv2 with FAISS significantly outperformed BEiTV2 (*χ*^2^ = 385.02, *p <* 0.001): of 496 discordant cases, DINOv2 was correct on 467 and BEiTV2 on 29, an odds ratio of 16.10 in DI-NOv2’s favour. DINOv2 also significantly outperformed BioCLIP (*χ*^2^ = 363.21, *p <* 0.001), winning 465 of 502 discordant cases, equivalent to DINOv2 being correct approximately 12.6 times as often as BioCLIP on disagreements. The BEiTV2 versus BioCLIP comparison did not reach statistical significance (*χ*^2^ = 0.18, *p* = 0.676): of 464 discordant cases, BEiTV2 was correct on 227 and BioCLIP on 237. Despite the 0.68 percentage point accuracy difference favouring BioCLIP (45.84% vs 45.16%), the supervised fine-tuned model and the true zero-shot model are statistically indistinguishable on the Kruger test set.

### Experiment 4 — Serengeti to Local Botswana Images

Table 11 presents the performance of all three models on the locally collected Botswana wildlife photography dataset (*n* = 316 images, 6 species). Unlike the Snapshot Kgalagadi and Snapshot Kruger test sets which were collected using standardised camera trap equipment, the local Botswana images are standard wildlife photographs captured by the research team using handheld cameras during daylight hours. The Maximum Mean Discrepancy between Serengeti and Botswana, computed using DINOv2 ViT-Large CLS token embeddings with an RBF kernel and median heuristic bandwidth (*σ* = 1.3002), was MMD^2^ = 0.0975, computed over *n* = 1,000 Serengeti and *m* = 316 Botswana images. This is the largest distributional distance of the three target datasets.

**Table 11.** Experiment 4: cross-domain performance on locally collected Botswana wildlife photographs (*n* = 316 images, 6 species: elephant, baboon, impala, hippo, giraffe, buffalo). Accuracy is reported with Wilson 95% confidence intervals. Performance drop in percentage points (pp) is measured relative to Experiment 1 accuracy. Macro F1 is computed over the 6 species present in this dataset only.

| Model | Accuracy % (95% CI) | Macro F1 (%) | Acc. Drop (pp) |
| --- | --- | --- | --- |
| DINOv2 + FAISS | 95.57 [92.70, 97.34] | 93.06 | +5.05 |
| BioCLIP | 92.72 [89.32, 95.10] | 90.93 | +45.96 |
| BEiT2 | 52.22 [46.71, 57.66] | 48.62 | −31.40 |

DINOv2 with FAISS achieved 95.57% accuracy (95% CI: 92.70–97.34%) and 93.06% macro F1, its highest performance across all four test sets and 5.05 percentage points above its in-domain Serengeti score of 90.52%. BioCLIP achieved 92.72% accuracy (95% CI: 89.32–95.10%) and 90.93% macro F1, also substantially above its performance on Kgalagadi (61.64%) and Kruger (45.84%).

DINOv2 achieved its highest accuracy on the dataset with the largest MMD distance from Serengeti, while it achieved its lowest accuracy on Kruger (MMD^2^ = 0.0348), the dataset with the smallest MMD distance. Feature-space distributional distance therefore did not predict cross-domain classification accuracy in this study.

BEiTV2 dropped to 52.22% accuracy (95% CI: 46.71–57.66%) and 48.62% macro F1 on the local Botswana images, a fall of 31.40 percentage points from its in-domain Serengeti baseline. It should be noted that the local Botswana images are standard wildlife photographs rather than camera trap sequences, and the same individual animal may appear across multiple images in this dataset. This introduces a potential source of dependency between test images that does not exist in the camera trap test sets.

Pairwise McNemar’s tests on Botswana yield a mixed pattern: BEiTV2 is significantly outperformed by both zero-shot models, but DINOv2 and BioCLIP are not statistically distinguishable from each other (Table 12).

**Table 12.** Experiment 4: pairwise McNemar’s test results on the local Botswana wildlife photography dataset.

| Model A | Model B | $n_{01}$ | $n_{10}$ | $\chi^2$ | $p$-value | OR |
| --- | --- | --- | --- | --- | --- | --- |
| BEiTV2 | DINOv2 + FAISS | 137 | 0 | 135.01 | $< 0.001$ | — |
| BEiTV2 | BioCLIP | 132 | 4 | 118.60 | $< 0.001$ | 33.00 |
| DINOv2 + FAISS | BioCLIP | 7 | 16 | 2.78 | 0.095 | 0.44 |

DINOv2 with FAISS significantly outperformed BEiTV2 (*χ*^2^ = 135.01, *p <* 0.001): of 137 discordant cases, DINOv2 was correct on all 137 and BEiTV2 on none, and the odds ratio is therefore undefined. BioCLIP also significantly outperformed BEiTV2 (*χ*^2^ = 118.60, *p <* 0.001), winning 132 of 136 discordant cases. The DINOv2 versus BioCLIP comparison did not reach statistical significance (*χ*^2^ = 2.78, *p* = 0.095): of 23 discordant cases, DINOv2 was correct on 16 and BioCLIP on 7. Despite the 2.85 percentage point accuracy gap (DINOv2 95.57% vs BioCLIP 92.72%), the small number of discordant cases on this 316-image test set does not provide sufficient statistical evidence to distinguish the two zero-shot models.

### Experiment 5 — Data Scaling Analysis

Table 13 presents the performance of BEiTV2 trained on four fractional subsets of the Snapshot Serengeti training split (1%, 5%, 25%, 50%) plus the full 100% checkpoint reused from Experiment 1, evaluated on all four test sets. Stratified subsets were generated with a single random seed across all fractions to guarantee nesting (the 1% subset is a subset of the 5% subset, the 5% subset of the 25% subset, and the 25% subset of the 50% subset). Training hyperparameters were held identical across all fractions and matched the Experiment 1 BEiTV2 configuration (Table 2). All four fractional trainings terminated by early stopping rather than reaching the maximum of 50 epochs.

**Table 13.**
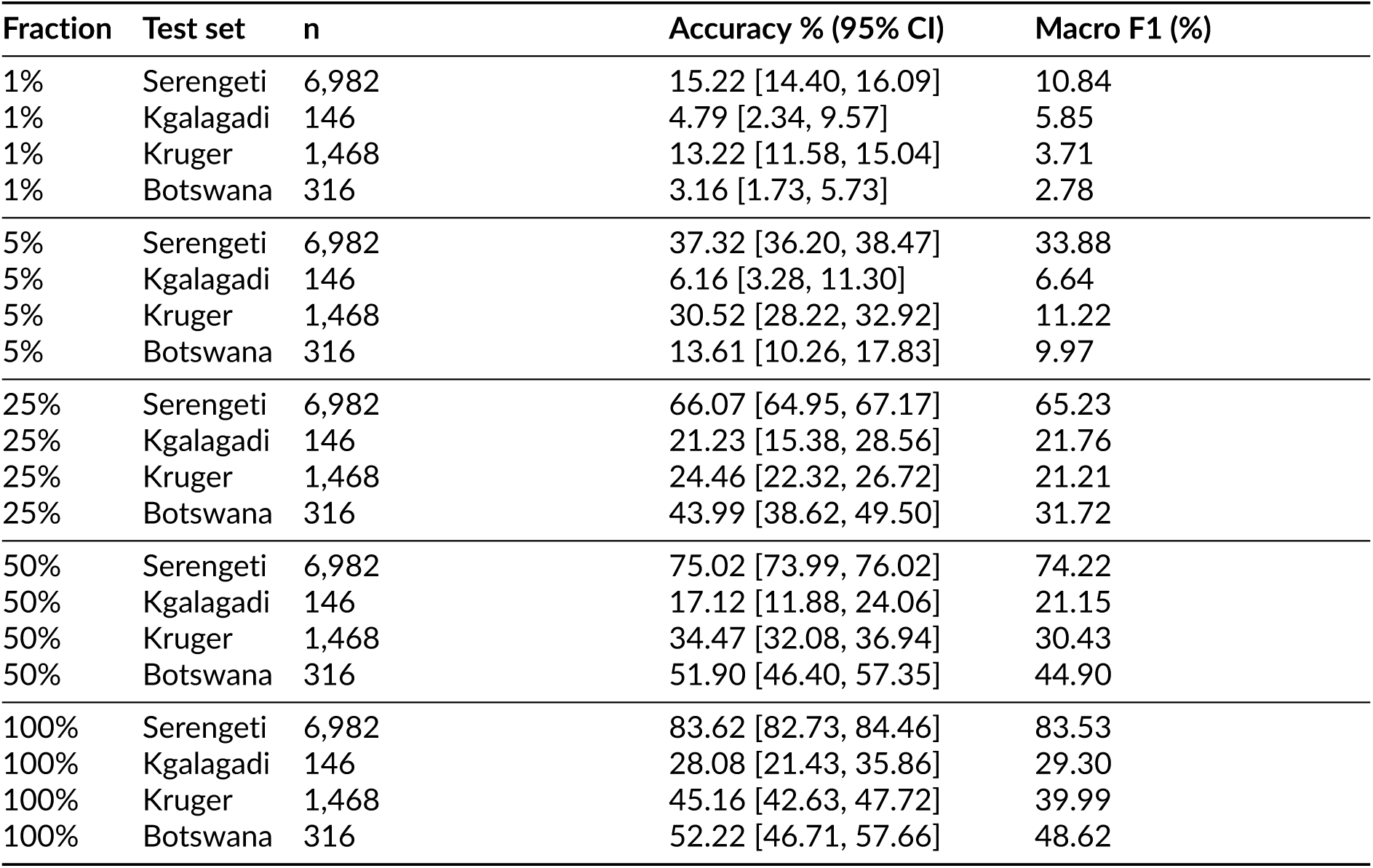
Experiment 5: full evaluation of all five BEiTV2 checkpoints on all four test sets under Convention A. Accuracy is reported with Wilson 95% confidence intervals. Macro F1 is computed over the classes present in each test set.

| Fraction | Test set | n | Accuracy % (95% CI) | Macro F1 (%) |
| --- | --- | --- | --- | --- |
| 1% | Serengeti | 6,982 | 15.22 [14.40, 16.09] | 10.84 |
| 1% | Kgalagadi | 146 | 4.79 [2.34, 9.57] | 5.85 |
| 1% | Kruger | 1,468 | 13.22 [11.58, 15.04] | 3.71 |
| 1% | Botswana | 316 | 3.16 [1.73, 5.73] | 2.78 |
| 5% | Serengeti | 6,982 | 37.32 [36.20, 38.47] | 33.88 |
| 5% | Kgalagadi | 146 | 6.16 [3.28, 11.30] | 6.64 |
| 5% | Kruger | 1,468 | 30.52 [28.22, 32.92] | 11.22 |
| 5% | Botswana | 316 | 13.61 [10.26, 17.83] | 9.97 |
| 25% | Serengeti | 6,982 | 66.07 [64.95, 67.17] | 65.23 |
| 25% | Kgalagadi | 146 | 21.23 [15.38, 28.56] | 21.76 |
| 25% | Kruger | 1,468 | 24.46 [22.32, 26.72] | 21.21 |
| 25% | Botswana | 316 | 43.99 [38.62, 49.50] | 31.72 |
| 50% | Serengeti | 6,982 | 75.02 [73.99, 76.02] | 74.22 |
| 50% | Kgalagadi | 146 | 17.12 [11.88, 24.06] | 21.15 |
| 50% | Kruger | 1,468 | 34.47 [32.08, 36.94] | 30.43 |
| 50% | Botswana | 316 | 51.90 [46.40, 57.35] | 44.90 |
| 100% | Serengeti | 6,982 | 83.62 [82.73, 84.46] | 83.53 |
| 100% | Kgalagadi | 146 | 28.08 [21.43, 35.86] | 29.30 |
| 100% | Kruger | 1,468 | 45.16 [42.63, 47.72] | 39.99 |
| 100% | Botswana | 316 | 52.22 [46.71, 57.66] | 48.62 |

On the in-domain Snapshot Serengeti test set, both accuracy and macro F1 increase strictly monotonically with training set size and exhibit clear diminishing returns (Figure 15). Accuracy rises from 15.22% at 1% data to 37.32% at 5%, 66.07% at 25%, 75.02% at 50%, and 83.62% at 100%. The 25% checkpoint reaches approximately 79% of the full-data accuracy on the in-domain test set, consistent with the diminishing-returns saturation behaviour reported by Bevan et al. (2026) for camera trap classification on the Maasai Mara and Bardia datasets above approximately 25% of available training data.

**Figure 15.**
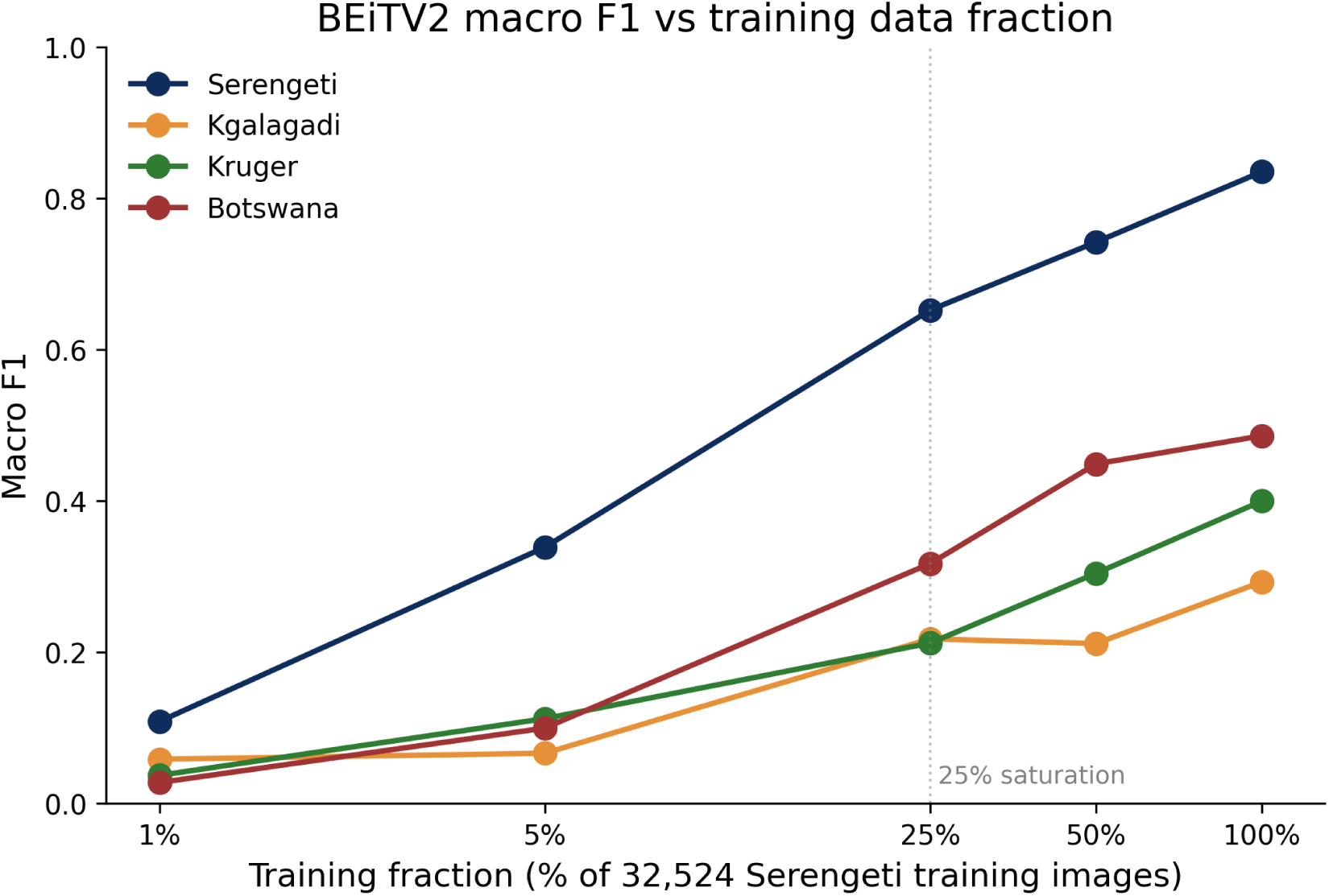
Experiment 5 macro F1 as a function of BEiTV2 training data fraction across all four test sets. The training fraction axis is log-scaled, with values 1%, 5%, 25%, 50%, and 100% of the 32,524-image Serengeti training split. The vertical reference line at 25% marks the saturation point identified from in-domain validation accuracy. In-domain Serengeti macro F1 rises steeply and exhibits clear diminishing returns above 25%. Cross-domain macro F1 also increases monotonically with training data, but with substantially weaker gains than in-domain across all three target test sets.

Cross-domain macro F1 also increases monotonically with training data on all three target test sets, but with substantially weaker gains than in-domain (Figure 15). Going from 25% to 100% data improves in-domain macro F1 by 18.30 percentage points (65.23% to 83.53%), but improves macro F1 by only 7.54 percentage points on Kgalagadi (21.76% to 29.30%), 18.78 percentage points on Kruger (21.21% to 39.99%), and 16.90 percentage points on Botswana (31.72% to 48.62%). The Kgalagadi result shows a small reversal between the 25% and 50% checkpoints (21.76% vs 21.15% macro F1), but the 95% confidence intervals on accuracy overlap substantially at this test set size (*n* = 146) and the difference falls within the noise floor implied by that sample size.

When measured by raw accuracy rather than macro F1, the cross-domain scaling pattern displays an apparent non-monotonicity on the Kruger test set: the 5% checkpoint achieves 30.52% accuracy ([28.22, 32.92]) while the 25% checkpoint achieves only 24.46% ([22.32, 26.72]). The two confidence intervals do not overlap, indicating that this accuracy reversal is not attributable to sampling noise. However, the macro F1 values for the same two checkpoints are 11.22% and 21.21% respectively, meaning that the 25% checkpoint achieves nearly twice the per-class average performance of the 5% checkpoint. Inspection of the per-class prediction distributions reveals the mechanism. The Kruger test set is heavily imbalanced, with impala constituting 585 of 1,468 test images (approximately 40%). The 5% checkpoint predicts impala on 798 of 1,468 Kruger images (54%), correctly classifying 363 of the 585 true impala images and defaulting to “impala” on a substantial fraction of images belonging to other species. This majority-class collapse inflates raw accuracy by capitalising on the dominance of impala in the test set distribution. The 25% checkpoint instead over-predicts dik-dik (506 dik-dik predictions for 19 true dik-dik images) and distributes its predictions more evenly across the 14 Kruger species, correctly classifying fewer impala (107) but more zebra, buffalo, giraffe, hare, hippo, and dik-dik than the 5% checkpoint. This more even prediction distribution yields a substantially higher macro F1 despite the lower aggregate accuracy.

The Botswana test set displays a distinct saturation pattern: macro F1 reaches 44.90% at 50% training data and improves only marginally to 48.62% at 100% training data, an increment of 3.72 percentage points from doubling the training set. Accuracy shows a similar plateau, rising only from 51.90% to 52.22%. This contrasts with both Serengeti (which gains 9.31 percentage points in macro F1 from 50% to 100%) and Kruger (which gains 9.56 percentage points). The Botswana test set differs from the camera trap datasets in image format, viewing angle, capture equipment, and lighting conditions, all of which are factors identified by Beery et al. (2018) as sources of location-specific overfitting. The apparent saturation of BEiTV2 performance on Botswana above 50% training data suggests that additional Serengeti camera trap data does not address the image-format-level domain shift represented by the locally collected wildlife photographs.

Across all four test sets, macro F1 increases monotonically with training data within statistical noise, while the cross-domain scaling is substantially weaker than the in-domain scaling. The Kruger accuracy reversal between the 5% and 25% checkpoints illustrates that on heavily imbalanced cross-domain test sets, aggregate accuracy can be inflated by majority-class artefacts that do not reflect genuine generalisation across species, supporting the adoption of macro F1 as the primary evaluation metric in this study.

### Experiment 6 — MegaDetector Preprocessing

Table 14 presents the performance of all three models evaluated on MegaDetector-cropped versions of all four test sets, alongside their raw-image baselines from Experiments 1–4. MegaDetector v6 with a YOLOv9c backbone (PytorchWildlife 1.1.0) was applied at a confidence threshold of 0.2 with 10% bounding box padding and square-padding to preserve aspect ratio, producing JPEG-90 crops. Detection rates were 94.6% on the Serengeti test set, 93.2% on Kgalagadi, 94.5% on Kruger, and 89.9% on Botswana; images for which no detection was found used the full uncropped image as a fallback. BEiTV2 was retrained from scratch on 32,522 MegaDetector-cropped Serengeti training images using the identical hyperparameter configuration as Experiment 1 (Table 2), converging after 33 epochs by early stopping with a validation accuracy of 87.58%. The DINOv2 FAISS retrieval index was rebuilt from cropped training image embeddings (32,522 vectors, 1024-dimensional, L2-normalised). BioCLIP inference was applied to cropped test images without any retraining, consistent with its zero-shot setup in Experiments 1–4.

**Table 14.** Experiment 6: classification performance on raw versus MegaDetector-cropped images for all three models across all four test sets. Accuracy is reported with Wilson 95% confidence intervals. The delta column is cropped minus raw accuracy in percentage points (pp). *χ*^2^ and *p*-value are from McNemar’s test with Edwards’ continuity correction comparing raw and cropped predictions on the same images; significance at *α* = 0.05 is marked with *.

| Test set | Model | n | Raw Acc<br>% (95%<br>CI) | Crop Acc<br>% (95%<br>CI) | $\Delta$ (pp) | Raw F1 | Crop F1 |
| --- | --- | --- | --- | --- | --- | --- | --- |
| Serengeti | BEiTV2 | 6,982 | 83.62<br>[82.73, 84.46] | 87.24<br>[86.41, 88.02] | +3.62* | 83.53 | 87.30 |
| Serengeti | DINOv2 + FAISS | 6,982 | 90.52<br>[89.81, 91.18] | 92.25<br>[91.62, 92.84] | +1.73* | 90.65 | 92.24 |
| Serengeti | BioCLIP | 6,982 | 46.76<br>[45.59, 47.93] | 58.75<br>[57.59, 59.90] | +11.99* | 47.53 | 58.96 |
| Kgalagadi | BEiTV2 | 146 | 28.08<br>[21.43, 35.86] | 64.38<br>[56.17, 71.83] | +36.30* | 29.30 | 69.36 |
| Kgalagadi | DINOv2 + FAISS | 146 | 77.40<br>[69.96, 83.43] | 86.99<br>[80.53, 91.59] | +9.59* | 79.81 | 87.95 |
| Kgalagadi | BioCLIP | 146 | 61.64<br>[53.55, 69.14] | 65.75<br>[57.65, 72.98] | +4.11 | 59.62 | 69.00 |
| Kruger | BEiTV2 | 1,468 | 45.16<br>[42.63, 47.72] | 55.45<br>[52.89, 57.99] | +10.29* | 39.99 | 54.34 |
| Kruger | DINOv2 + FAISS | 1,468 | 75.00<br>[72.72, 77.15] | 80.11<br>[77.99, 82.07] | +5.11* | 71.69 | 74.22 |
| Kruger | BioCLIP | 1,468 | 45.84<br>[43.31, 48.40] | 57.08<br>[54.53, 59.59] | +11.24* | 44.79 | 46.62 |
| Botswana | BEiTV2 | 316 | 52.22<br>[46.71, 57.66] | 69.94<br>[64.63, 74.73] | +17.72* | 48.62 | 64.96 |
| Botswana | DINOv2 + FAISS | 316 | 95.57<br>[92.70, 97.34] | 95.25<br>[92.30, 97.12] | −0.32 | 93.06 | 95.85 |
| Botswana | BioCLIP | 316 | 92.72<br>[89.32, 95.10] | 93.35<br>[90.06, 95.60] | +0.63 | 90.93 | 91.72 |

MegaDetector cropping improved classification accuracy for 11 of the 12 (test set, model) pairs, with 9 of the 12 improvements statistically significant at *α* = 0.05 by McNemar’s test. The three non-significant cases are Kgalagadi BioCLIP (+4.11 pp, *p* = 0.377), Botswana DINOv2 (*−*0.32 pp, *p* = 1.000), and Botswana BioCLIP (+0.63 pp, *p* = 0.838).

The largest benefit of cropping accrues to BEiTV2, the supervised model, across all four test sets. On the in-domain Serengeti test set, cropping raises BEiTV2 accuracy from 83.62% to 87.24% (+3.62 pp) and macro F1 from 83.53% to 87.30%. The cross-domain gains are substantially larger: +36.30 pp on Kgalagadi (28.08% to 64.38%), +10.29 pp on Kruger (45.16% to 55.45%), and +17.72 pp on Botswana (52.22% to 69.94%). The macro F1 gains mirror the accuracy gains; on Kgalagadi, BEiTV2 macro F1 rises from 29.30% to 69.36%, nearly a 40-percentage-point improvement from providing the model with a cropped animal region rather than the full background-containing frame. These results suggest that a substantial fraction of BEiTV2’s cross-domain degradation reported in Experiments 2–4 is attributable to background and contextual cues that differ between Serengeti and the target ecosystems, rather than to differences in the appearance of the animal species themselves.

DINOv2 with FAISS also benefits from cropping on the camera trap test sets, with statistically significant gains of +1.73 pp on Serengeti, +9.59 pp on Kgalagadi, and +5.11 pp on Kruger. The Botswana result is an exception: cropped DINOv2 accuracy falls marginally from 95.57% to 95.25% (*−*0.32 pp, *p* = 1.000), a difference of a single image on a 316-image test set. The Botswana dataset already approaches the performance ceiling for this task under raw-image conditions, and the detection-rate fallback of 10.1% (the lowest among the four test sets) means that approximately 32 Botswana images are evaluated on full uncropped frames. For the Botswana set it is also worth noting that macro F1 *increases* from 93.06% to 95.85% with cropping even as accuracy falls marginally, indicating that the single misclassification introduced by cropping involves a common class, not a structurally worse classification pattern. Overall, cropping improves DINOv2’s retrieval-index architecture by reducing the influence of location-specific background features that are encoded into the DINOv2 embedding even without supervised fine-tuning.

BioCLIP shows the most heterogeneous response to cropping. The gains on Serengeti (+11.99 pp) and Kruger (+11.24 pp) are among the largest in the table for their respective test sets, despite BioCLIP being a zero-shot model that receives no information about the training distribution. The Kgalagadi gain (+4.11 pp) does not reach statistical significance, consistent with the small test set size (*n* = 146). On Botswana, where BioCLIP already achieves 92.72% under raw-image conditions, the +0.63 pp gain is negligible and non-significant. The macro F1 pattern for BioCLIP is less consistent than for BEiTV2 and DINOv2: Kruger BioCLIP macro F1 rises only modestly from 44.79% to 46.62% despite an 11.24 pp accuracy gain, suggesting that the accuracy improvement on Kruger reflects gains concentrated on the dominant impala class rather than balanced per-species improvement.

Compared to the relative error reductions reported by Vyskočil and Picek (2025) for MegaDetector cropping on American and European camera trap datasets (42% on CCT20, 48% on CEF22, 75% on WCT), the gains for BEiTV2 on Kgalagadi and Botswana are of comparable magnitude in absolute accuracy terms, though direct comparison is complicated by differences in dataset size, number of classes, and the fact that Vyskočil and Picek (2025) employed a two-stage pipeline (a separate classifier for cropped and uncropped images) while we retrain a single classifier on cropped images only. The consistently positive direction of the effect across all three models and all four test sets — including the zero-shot models BioCLIP and DINOv2 that were not retrained on cropped images — supports the conclusion that the benefit of MegaDetector cropping is not an artefact of retraining but reflects a genuine reduction in the influence of background context on embedding quality across multiple architectures.

### Experiment 7 — Grayscale vs Colour

Table 15 presents the performance of all three models on the four test sets under both colour and grayscale conditions (*n* = 6,982 for Serengeti, 146 for Kgalagadi, 1,468 for Kruger, 316 for Botswana). Grayscale was applied dynamically at evaluation time by converting each test image to single-channel luminance and replicating it across the three RGB channels before applying the standard Convention A transform. For DINOv2 with FAISS, only the test queries were converted to grayscale; the retrieval index was retained in its original colour-feature form, mirroring the realistic deployment scenario in which a curated colour database is queried by infrared or low-light images collected at deployment sites. Pairwise McNemar’s tests with Edwards’ continuity correction were applied to compare colour and grayscale predictions on the same images for each (test set, model) pair.

**Table 15.** Experiment 7: classification performance under colour and grayscale conditions for all three models across all four test sets. Accuracy is reported with Wilson 95% confidence intervals. The accuracy drop in percentage points (pp) is colour minus grayscale. *χ*^2^, *p*-value, and the significance verdict at *α* = 0.05 are from McNemar’s test on the paired predictions.

| Test set | Model | Acc. col. % | Acc. gray % | Drop (pp) | $\chi^2$ | $p$-value |
| --- | --- | --- | --- | --- | --- | --- |
| Serengeti | BEiTV2 | 83.62 | 64.87 | 18.75 | 999.92 | < 0.001* |
| Serengeti | DINOv2 | 90.52 | 87.60 | 2.92 | 98.59 | < 0.001* |
| Serengeti | BioCLIP | 46.76 | 41.19 | 5.57 | 106.39 | < 0.001* |
| Kgalagadi | BEiTV2 | 28.08 | 19.18 | 8.90 | 5.33 | 0.0209* |
| Kgalagadi | DINOv2 | 77.40 | 78.08 | −0.68 | 0.00 | 1.0000 |
| Kgalagadi | BioCLIP | 61.64 | 56.85 | 4.79 | 1.03 | 0.3105 |
| Kruger | BEiTV2 | 45.16 | 25.14 | 20.03 | 255.50 | < 0.001* |
| Kruger | DINOv2 | 75.00 | 73.09 | 1.91 | 10.41 | 0.0013* |
| Kruger | BioCLIP | 45.84 | 33.45 | 12.40 | 126.98 | < 0.001* |
| Botswana | BEiTV2 | 52.22 | 37.34 | 14.87 | 23.25 | < 0.001* |
| Botswana | DINOv2 | 95.57 | 93.67 | 1.90 | 4.17 | 0.0412* |
| Botswana | BioCLIP | 92.72 | 82.91 | 9.81 | 25.71 | < 0.001* |

DINOv2 with FAISS was the most colour-robust of the three models across all four test sets, with accuracy drops ranging from *−*0.68 to 2.92 percentage points. On Kgalagadi, the grayscale accuracy of 78.08% slightly exceeded the colour accuracy of 77.40%, a difference of one correctly classified image out of 146 that McNemar’s test confirms is statistically indistinguishable from zero (*χ*^2^ = 0.00, *p*= 1.000). On the remaining three test sets, DINOv2’s grayscale accuracy was significantly lower than its colour accuracy at *α* = 0.05, but in absolute terms the drops were modest (2.92 pp on Serengeti, 1.91 pp on Kruger, 1.90 pp on Botswana). The self-supervised pretraining of DINOv2 on 142 million images appears to produce features that depend only weakly on colour information for the species classification task.

BEiTV2 was the most colour-dependent of the three models, with significant accuracy drops of 18.75 pp on Serengeti, 8.90 pp on Kgalagadi, 20.03 pp on Kruger, and 14.87 pp on Botswana. The drop on Kruger reduces BEiTV2’s accuracy from 45.16% to 25.14%, taking it well below the in-domain BEiTV2 colour accuracy on Serengeti even before any geographic domain shift is accounted for. This substantial degradation reflects BEiTV2’s reliance on colour-specific features learned during supervised fine-tuning on the predominantly colour Serengeti training set, which do not generalise to grayscale inputs.

BioCLIP occupied an intermediate position, with significant drops on Serengeti (5.57 pp), Kruger (12.40 pp), and Botswana (9.81 pp), but no statistically detectable drop on Kgalagadi (*χ*^2^ = 1.03, *p* = 0.3105). The non-significant Kgalagadi result is partly attributable to the small test set size (*n* = 146) and the already modest colour accuracy of 61.64%, which together constrain the McNemar test’s ability to detect a real difference; the 4.79 pp drop in raw accuracy is consistent in direction with BioCLIP’s behaviour on the other three test sets.

These results have direct implications for real-world camera trap deployment. Cameras frequently capture grayscale infrared images at night, particularly in ecosystems with high nocturnal activity. As reported in the Methodology section, Kruger has 35.3% nighttime captures based on timestamp metadata. The BEiTV2 grayscale performance on Kruger (25.14%) is closer to random chance for a 14-class problem (7.1%) than to its colour performance (45.16%), suggesting that supervised BEiTV2 would substantially underperform in deployment settings where night captures are common. DINOv2’s robustness across colour and grayscale conditions, by contrast, suggests it would maintain its accuracy advantage over BEiTV2 under such realistic deployment conditions.

### Experiment 8 — Per-Species Transfer Analysis

To identify which species suffer most from geographic domain shift and to characterise the dominant failure modes of each model, we computed per-species precision, recall, and F1 score for each of the three cross-domain test sets (Kgalagadi, Kruger, and Botswana) under each of the three models. Per-species metrics were derived directly from the prediction files of Experiments 2, 3, and 4, with no new inference performed. Confusion matrices were constructed in the canonical Serengeti 17-class index space, retaining the full matrix even for test sets containing fewer species, to support identification of misclassification patterns across the source taxonomy.

Tables 16–17 summarise the top per-species performers, the worst per-species performers, and the dominant species-to-species confusions for each cell of the 3 *×* 3 test set *×* model grid.

**Table 16.** Experiment 8: best and worst performing species (by F1) for each model on each cross-domain test set. Numbers in parentheses are the per-species F1 score.

| Test set | Model | Best species (F1) | Worst species (F1) |
| --- | --- | --- | --- |
| Kgalagadi | BEiTV2 | ostrich (0.65) | jackal (0.00) |
| Kgalagadi | DINOv2 | ostrich (0.86), eland (0.88) | wildebeest (0.69) |
| Kgalagadi | BioCLIP | ostrich (0.77) | wildebeest (0.27) |
| Kruger | BEiTV2 | elephant (0.51) | dik-dik (0.07), waterbuck (0.00) |
| Kruger | DINOv2 | elephant (0.76), zebra (0.82) | dik-dik (0.14), waterbuck (0.00) |
| Kruger | BioCLIP | impala (0.49), kori bustard – | dik-dik (0.12), waterbuck (0.00) |
| Botswana | BEiTV2 | baboon (0.79) | buffalo (0.11) |
| Botswana | DINOv2 | baboon (1.00), elephant (0.96) | buffalo (0.67) |
| Botswana | BioCLIP | elephant (0.97) | buffalo (0.61) |

**Table 17.** Experiment 8: dominant species-to-species confusions per test set and model, ranked by absolute count. The "True *→* Predicted" column shows the misclassification direction; "Count" is the number of test images where this confusion occurred.

| Test set | Model | True → Predicted | Count |
| --- | --- | --- | --- |
| Kruger | BEiTV2 | impala → dik-dik | 208 |
| Kruger | DINOv2 | impala → dik-dik | 162 |
| Kruger | BioCLIP | impala → kori bustard | 122 |
| Kruger | BioCLIP | impala → dik-dik | 96 |
| Botswana | BEiTV2 | elephant → buffalo | 46 |

A consistent pattern emerges across the three cross-domain test sets: the species that are hardest to classify under domain shift are those with small representation in the target test set (dik-dik with 19 Kruger images, waterbuck with 43 Kruger images, buffalo with 11 Botswana images), and the species that are easiest are those with distinctive visual features and adequate representation (ostrich, elephant, zebra, baboon). DINOv2 with FAISS achieves an F1 of 1.00 on Botswana baboons, correctly classifying all 52 baboon images with zero false positives, the strongest per-species result anywhere in the cross-domain analysis.

### Dominant Cross-Domain Confusions

Table 17 reports the most frequent species-to-species mis-classifications for each cell, ranked by the absolute number of confused images.

The impala *→* dik-dik confusion is the dominant failure mode on Kruger across all three models, although it manifests with different severity. BEiTV2 misclassifies 208 of 585 Kruger impala images as dik-dik (35.6%), and DINOv2 misclassifies 162 (27.7%) despite its substantially higher overall recall on impala. BioCLIP’s top confusion is instead impala *→* kori bustard (122 images), with impala *→* dik-dik in second place (96 images). The dik-dik over-prediction is striking: Kruger contains only 19 true dik-dik images, yet BEiTV2 predicts dik-dik 333 times (321 false positives), DINOv2 predicts dik-dik 222 times (205 false positives), and BioCLIP predicts dik-dik 193 times (180 false positives). All three models substantially over-predict dik-dik on Kruger imagery despite its near-absence from the test distribution, suggesting that the feature-space embeddings of medium-sized antelopes in Kruger camera trap photography align more closely with the dik-dik features learned from Serengeti than with the impala features, even though the model was trained on far more Serengeti impala (*n*_train_= 2,099) than dik-dik (*n*_train_ = 2,100).

On Botswana, BEiTV2 misclassifies 46 of 177 elephant images as buffalo (26.0%), which is the single dominant failure mode driving BEiTV2’s weak Botswana performance. The resulting buffalo F1 of 0.108 reflects a combination of low recall (4 of 11 true buffalo correctly identified) and high false positive count (59 false buffalo predictions, of which 46 are misclassified elephants). This elephant *→* buffalo bias is unique to BEiTV2 on Botswana; DINOv2 and BioCLIP do not exhibit it, with elephant F1 scores of 0.96 and 0.97 respectively on the same test set.

#### Effect of Training Data Volume on Per-Species Performanc

Per-species F1 on the cross-domain test sets shows no clear relationship with the number of Serengeti training images per species. The Serengeti training split contains between 700 and 2,100 images per species. Hare and waterbuck, the two species with the fewest training images (700 each), are not consistently the worst performers cross-domain: hare achieves the second-highest recall on Kruger BEiTV2 (after impala), while waterbuck achieves F1 of 0.00 under all three models on Kruger. Conversely, dik-dik (2,100 training images, near the top of the training distribution) is consistently the worst performer on Kruger across all three models. The dominant determinant of per-species cross-domain performance therefore appears to be visual similarity to confusable species in the target test set, rather than the number of training images available for the species in the source domain.

## Discussion

### Geographic Domain Shift Affects Models Differently

The results presented in Experiments 2–4 reveal that the three evaluated models respond very differently to geographic domain shift from the East African Serengeti training data to the three African target regions. BEiTV2, the supervised fine-tuned model, suffered the most severe degradation: accuracy fell by 55.54 percentage points on Kgalagadi, 38.46 percentage points on Kruger, and 31.40 percentage points on Botswana relative to the in-domain Serengeti baseline. DINOv2 with FAISS retained the strongest cross-domain performance, with accuracy drops of only 13.12 percentage points on Kgalagadi and 15.52 percentage points on Kruger, and an accuracy increase of 5.05 percentage points on Botswana. BioCLIP showed the most variable behaviour, with apparent accuracy increases on Kgalagadi (+14.88 pp) and Botswana (+45.96 pp), but a smaller accuracy change on Kruger (*−*0.92 pp); these increases partly reflect the smaller class spaces of the target test sets (5 species on Kgalagadi and 6 on Botswana versus 17 on Serengeti) rather than improved species discrimination.

The contrast between BEiTV2 and DINOv2 is methodologically informative. Both models share a vision transformer backbone of similar capacity, but they differ in their training paradigm: BEiTV2 is supervised and learns to discriminate the 17 Serengeti species, while DINOv2 is self-supervised on a curated 142 million-image dataset and produces generic visual features unconstrained by species-classification objectives. The substantially smaller cross-domain accuracy drop for DINOv2 suggests that supervised fine-tuning on a single ecosystem encodes location-specific or species-arrangement-specific cues that do not transfer to new geographic regions, supporting the location-overfitting hypothesis raised by Beery et al. (2018).

### Distributional Distance Does Not Predict Cross-Domain Accuracy

We computed Maximum Mean Discrepancy (MMD) between DINOv2 embeddings of the source and each target dataset under an RBF kernel with median heuristic bandwidth. The squared MMD values were MMD^2^ = 0.0348 for Serengeti–Kruger, 0.0706 for Serengeti–Kgalagadi, and 0.0975 for Serengeti–Botswana. If MMD in DINOv2 feature space were a reliable proxy for the difficulty of cross-domain transfer, we would expect classification accuracy to correlate inversely with MMD. Instead, DINOv2 achieved its highest accuracy (95.57%) on Botswana (largest MMD) and its lowest accuracy (75.00%) on Kruger (smallest MMD). For BEiTV2 the pattern is similarly uncorrelated: accuracy is highest on Botswana (52.22%, largest MMD) and lowest on Kgalagadi (28.08%, middle MMD).

This finding has practical implications. MMD and similar feature-space distance metrics are widely used as proxies for domain gap severity (Gretton et al., 2012), but our results indicate that for camera trap species classification across African ecosystems, the distance computed in a generic pretrained feature space does not predict transfer performance. Other factors — image format, viewing angle, lighting conditions, animal pose distribution, and species co-occurrence patterns — appear to drive the cross-domain accuracy variation more strongly than overall distributional divergence. Conservation practitioners should be cautious about relying on MMD-style domain-distance metrics when planning cross-region model deployment.

### Supervised Training Offers No Advantage on Kruger

A striking finding from Experiment 3 is that BEiTV2 and BioCLIP achieved statistically indistinguishable accuracy on Kruger (45.16% vs 45.84%; McNemar’s *χ*^2^ = 0.18, *p* = 0.676). BEiTV2 was fine-tuned on 32,524 labelled Serengeti images while BioCLIP received zero training data from any of our datasets. The fact that supervised training on 32,524 in-region images provides no measurable advantage over pure zero-shot inference on this test set challenges the assumption that labelled training data is always preferable to zero-shot foundation models for cross-region deployment.

The likely explanation is that the supervised model’s discriminative features overfit to Serengeti-specific image characteristics — camera placement, viewing angle, lighting, animal context — that do not transfer cleanly to Kruger’s different camera trap deployment style. BioCLIP, having learned diverse biological imagery from curated taxonomic sources, brings broader visual priors that work approximately as well as the supervised features on this particular cross-domain task. This finding suggests that the cost-benefit calculation for camera trap classification in conservation settings should not assume supervised fine-tuning is worth the annotation effort, particularly when the target deployment region differs visually from the available labelled training data.

### Image-Format Domain Shift Imposes a Ceiling

The Botswana results in Experiments 4, 5, and 7 collectively suggest that some domain shifts cannot be addressed by additional in-domain training data. The local Botswana dataset consists of handheld wildlife photographs rather than camera trap sequences — a fundamentally different image format from the Serengeti training data. BEiTV2, trained exclusively on camera trap images, achieved 52.22% accuracy on Botswana with the full training set. Experiment 5 showed that this accuracy improved by only 0.32 percentage points from the 50% training fraction (51.90%) to 100%. The dominant failure mode is BEiTV2 misclassifying Botswana elephants as buffalo (46 of 177 elephant images; Experiment 8), a confusion that does not appear in DI-NOv2 (F1 of 0.96 on Botswana elephants) or BioCLIP (F1 of 0.97).

This pattern is consistent with the location-specific overfitting hypothesis extended to image-format-specific overfitting: a model trained on camera-trap-style imagery learns features that depend on the camera trap framing, scale, lighting profile, and overlay artefacts, and these features do not generalise to handheld photographic imagery even when the species concept is unchanged. The implication for practitioners is that an additional axis of domain shift — image-format mismatch — can impose a performance ceiling that more in-format training data cannot address. Models intended for diverse-format deployment may require either training data that spans multiple capture formats, or reliance on pretrained-from-diverse-data foundation models such as DINOv2 which include handheld photographic imagery in their pretraining distribution.

### Data Scaling Is Substantially Weaker Across Domains

Experiment 5 reveals a clear difference between in-domain and cross-domain data scaling. In-domain Serengeti macro F1 increases strictly monotonically with training data from 1% (0.1084) to 100% (0.8353), reproducing the saturation pattern documented by Bevan et al. (2026). Cross-domain macro F1 also increases monotonically with training data, but the gains are substantially weaker: going from 25% to 100% training data improves in-domain macro F1 by 18.30 percentage points but improves cross-domain macro F1 by only 7.54 percentage points on Kgalagadi, 18.78 on Kruger, and 16.90 on Botswana. Equivalently, the 25% checkpoint reaches 79% of full-data macro F1 on Serengeti but only 74% of full-data macro F1 on Kgalagadi.

The Experiment 5 results also illustrate a methodological point that generalises beyond data scaling. The Kruger test set exhibits an apparent non-monotonicity in raw accuracy between the 5% and 25% checkpoints (30.52% vs 24.46%), with non-overlapping 95% confidence intervals. Inspection of per-class predictions revealed that the 5% checkpoint defaults to predicting impala on 54% of Kruger images (vs the 40% true prevalence), inflating raw accuracy through majority-class collapse rather than genuine species discrimination. The 25% checkpoint distributes its predictions more evenly and achieves nearly double the macro F1 (0.1122 vs 0.2121) despite the lower aggregate accuracy. On heavily imbalanced cross-domain test sets, aggregate accuracy can be a misleading measure of model quality; per-class metrics such as macro F1 are necessary to distinguish models that have learned species- discriminative features from those that have learned only the target test distribution’s class prior.

### Grayscale and Colour Robustness

Experiment 7 evaluated all three models under grayscale conditions, simulating the infrared night captures that are common in camera trap deployments. DINOv2 with FAISS proved most colour-robust, with accuracy drops ranging from *−*0.68 to 2.92 percentage points across all four test sets. BEiTV2 was most colour-dependent, with drops of 18.75 to 20.03 percentage points on the three larger test sets, all statistically significant. BioCLIP occupied an intermediate position with significant drops on Serengeti, Kruger, and Botswana but no detectable drop on Kgalagadi (likely reflecting the small Kgalagadi test set size of 146 images).

The implication for real-world deployment is substantial. As reported in the Image Analysis section, Kruger has 35.3% nighttime captures based on timestamp metadata, with many of these returning grayscale infrared images. BEiTV2’s grayscale accuracy on Kruger of 25.14% is closer to random chance for a 14-class problem (7.1%) than to its colour accuracy of 45.16%. In deployment settings with substantial nighttime activity, BEiTV2 would substantially underperform its colour-only validation results. DINOv2’s robustness across colour and grayscale conditions makes it a better candidate for deployment in such settings.

### MegaDetector Preprocessing

MegaDetector cropping had a substantial and broadly consistent positive effect on classification accuracy across all three models in our study. For BEiTV2, cropping improved cross-domain accuracy by an average of 21.44 percentage points across the three target test sets (Kgalagadi +36.30 pp, Kruger +10.29 pp, Botswana +17.72 pp), all statistically significant, with the in-domain Serengeti improvement of +3.62 pp significant as well. For DINOv2 with FAISS, cropping yielded significant gains on the three camera trap test sets (Serengeti +1.73 pp, Kgalagadi +9.59 pp, Kruger +5.11 pp) with no meaningful change on Botswana where performance was already near ceiling. For BioCLIP, significant gains were observed on Serengeti (+11.99 pp) and Kruger (+11.24 pp), with smaller non-significant changes on Kgalagadi and Botswana.

The magnitude of the BEiTV2 benefit is particularly notable. The 36.30 pp accuracy gain on Kgalagadi — from 28.08% to 64.38% — substantially closes the gap between BEiTV2 and DINOv2 on that test set (from a 49.32 pp gap under raw conditions to a 22.61 pp gap under cropped conditions). This suggests that a large fraction of BEiTV2’s cross-domain degradation in Experiments 2–4 is driven by background and habitat context cues that change between Serengeti and the target ecosystems, rather than by appearance differences in the animal species themselves. When the model is forced to classify based on a tight crop around the detected animal, the habitat-context component of the domain gap is substantially reduced, and the species-discriminative features that BEiTV2 has learned transfer more successfully.

The fact that DINOv2 also benefits from cropping, despite not being retrained on cropped images (its index was rebuilt from cropped training features, but the underlying feature extractor is unchanged), indicates that background context influences the retrieval embedding even for a self-supervised model. The smaller absolute gains for DINOv2 compared to BEiTV2 are consistent with DINOv2’s self-supervised pretraining having already produced features that are less background- dependent than the species-specific features learned by BEiTV2’s supervised fine-tuning. Similarly, the BioCLIP gains on Serengeti and Kruger confirm that zero-shot text-embedding similarity is also affected by background context, even though BioCLIP was never trained on our datasets.

The three non-significant cases (Kgalagadi BioCLIP, Botswana DINOv2, Botswana BioCLIP) share a common explanation. For Botswana DINOv2 and BioCLIP, performance under raw conditions (95.57% and 92.72% respectively) leaves little room for improvement, and the 10.1% fallback rate on Botswana means approximately 32 images are processed without a crop. For Kgalagadi BioCLIP, the small test set size (*n* = 146) limits statistical power; the 4.11 pp raw accuracy gain is consistent in direction with BioCLIP’s behaviour on the other three test sets.

MegaDetector v6 achieved detection rates of 89.9–94.6% across our test sets. The reported cropped accuracies therefore reflect a mixture of cropped and full-image fallback inference, weighted by the per-test-set detection rate. The 10.1% fallback rate on Botswana is the highest, reflecting the greater variation in image composition in the handheld wildlife photography dataset compared to standardised camera trap images.

### Limitations

This study has several limitations worth acknowledging. First, the local Botswana dataset is small (*n* = 316) and consists of standard wildlife photographs rather than camera trap sequences. As noted in Experiment 4, the same individual animal may appear across multiple photographs, introducing a potential dependency between test images that does not exist in the camera trap test sets. The high accuracy of DINOv2 (95.57%) and BioCLIP (92.72%) on Botswana should be interpreted with this caveat in mind: the effective sample size of distinct animal individuals is smaller than the image count suggests, and confidence intervals would be wider under a per-individual rather than per-image analysis.

Second, the cross-domain test sets are small after applying the minimum-images-per-species threshold of nine: 146 images for Kgalagadi and 316 for Botswana. The resulting 95% confidence intervals on accuracy span 13 to 14 percentage points on these test sets, limiting the precision with which model differences can be estimated. The Kruger test set is larger (*n* = 1,468) and yields correspondingly tighter intervals.

Third, the supervised BEiTV2 model was fine-tuned only on Serengeti. We did not explore multi-region supervised training (e.g. training on a mixture of Serengeti, Kruger, and Kgalagadi labelled data), which might address some of the cross-domain failure modes identified here. Such an extension is left to future work.

Fourth, DINOv2 with FAISS retrieves nearest neighbours from a Serengeti-only image database. The retrieval-based approach allows trivial extension by adding labelled images from new regions to the database; we did not test this in our cross-domain experiments to keep the comparison with BEiTV2 fair (both models see only Serengeti labels), but in practice this is a meaningful advantage of the retrieval approach for conservation deployment.

### Implications for Conservation Practice

For conservation practitioners considering deployment of automated species classification across African ecosystems, our results suggest three practical recommendations.

First, retrieval-based zero-shot classification using DINOv2 embeddings appears to be the most robust general-purpose approach across the geographic shifts examined here. It avoids both the location-specific overfitting of supervised fine-tuning and the labelling and retraining cost of new region adaptation, while remaining substantially more accurate than the text-prompt zero-shot approach (BioCLIP) on most test sets.

Second, supervised fine-tuning of camera trap species classifiers should not be assumed to outperform zero-shot foundation models in cross-region deployment. On our Kruger test set, BEiTV2 and BioCLIP were statistically indistinguishable despite BEiTV2 receiving over 30,000 labelled training images and BioCLIP receiving none. Practitioners should consider the cost-benefit of annotation effort against the achievable cross-region accuracy gain.

Third, image-format mismatch (camera trap vs handheld photography, or daytime colour vs nighttime grayscale) represents a domain shift that more in-domain training data cannot fully address. Deployments in mixed-format or mixed-condition settings should plan around this limitation, either by including diverse-format training data or by choosing models with diverse-format pretraining (such as DINOv2 or BioCLIP).

## Conclusion

This study evaluated three contrasting approaches to wildlife species classification — supervised fine-tuning (BEiTV2), retrieval-based zero-shot (DINOv2 + FAISS), and text-prompt zero-shot (BioCLIP) — across the in-domain Snapshot Serengeti dataset and three African cross-domain test sets representing distinct ecosystems: Snapshot Kgalagadi, Snapshot Kruger, and a locally collected Botswana wildlife photography dataset. Eight experiments examined in-domain baselines, geographic transferability, training data scaling, MegaDetector preprocessing, colour robustness, and per-species transfer behaviour.

The results demonstrate that DINOv2 with FAISS, despite requiring no model training, achieves the strongest and most stable performance across all four test sets, with macro F1 scores of 90.65% (Serengeti in-domain), 79.81% (Kgalagadi), 71.69% (Kruger), and 93.06% (Botswana). The supervised BEiTV2 model exhibits substantially greater cross-domain degradation, including the striking finding that supervised fine-tuning on 32,524 Serengeti images provides no statistically significant advantage over zero-shot BioCLIP on the Kruger test set. Maximum Mean Discrepancy between datasets in DINOv2 feature space does not predict cross-domain accuracy in our results, suggesting that feature-space distributional metrics are insufficient proxies for transfer difficulty in this domain.

Several findings have direct practical relevance for conservation practitioners. Training data scaling shows substantially weaker gains on cross-domain test sets than on in-domain data, suggesting that more labelled data is not a complete remedy for geographic shift. BEiTV2 is highly colour-dependent, losing 18 to 20 percentage points of accuracy when test images are converted to grayscale; DINOv2 is comparatively colour-robust, losing at most 3 percentage points. On heavily imbalanced cross-domain test sets, aggregate accuracy can be inflated by majority-class artefacts; macro F1 is necessary to distinguish models that have learned species-discriminative features from those that have learned only the target test distribution’s class prior. Image-format domain shift (camera trap vs handheld photography) imposes a performance ceiling on BEiTV2 that more in-domain training cannot address.

MegaDetector preprocessing substantially improved classification performance across all three models and all four test sets, with statistically significant gains in 9 of 12 conditions; BEiTV2 benefited most, gaining an average of 21.44 percentage points in cross-domain accuracy, indicating that a substantial component of supervised model domain shift is attributable to background and habitat context rather than species appearance.

Future work should explore multi-region supervised training, deeper analysis of which species-level features transfer across ecosystems, and the practical viability of retrieval-based classification as a deployment paradigm for conservation applications across African wildlife monitoring programs.

## Acknowledgements

The authors acknowledge the assistance of Claude (Anthropic) as an AI writing assistant in improving the clarity, grammar, and quality of the manuscript, and in providing code assistance during software development. All scientific content, analyses, results, and interpretations were reviewed and verified by the authors.

## Data, script, code, and supplementary information availability

Camera trap datasets are publicly available through the Labeled Information Library of Alexandria (LILA BC) at https://lila.science. The Snapshot Serengeti, Snapshot Kgalagadi, and Snapshot Kruger datasets are all accessible through LILA BC. The local Botswana wildlife photography dataset, along with all analysis scripts and code for reproducing the experiments, is available on Zenodo at https://doi.org/10.5281/zenodo.20774512 (Nanduri et al., 2026).

## References

Beery S, Morris D, Perona P (2019a). The iWildCam 2019 Challenge Dataset. arXiv: 1907.07617 [cs.CV]. URL: https://arxiv.org/abs/1907.07617.

Beery S, Morris D, Yang S (2019b). Efficient pipeline for camera trap image review. 10.48550/arXiv.1907.06772.

Beery S, Van Horn G, Perona P (2018). Recognition in Terra Incognita. In: Proceedings of the European Conference on Computer Vision (ECCV), pp. 472–489. 10.1007/978-3-030-01270-0_28.

Bevan PA, Pantazis O, Pringle HA, Braga Ferreira G, Ingram DJ, Madsen EK, Thomas L, Thanet DR, Silwal T, Rayamajhi S, Brostow GJ, Mac Aodha O, Jones KE (2026). Deep learning-based ecological analysis of camera trap images is impacted by training data quality and quantity. Remote Sensing in Ecology and Conservation. 10.1002/rse2.70052.

Burton AC, Neilson E, Moreira-Arce D, Ladle A, Steenweg R, Fisher JT, Bayne E, Boutin S (2015). Wildlife camera trapping: a review and recommendations for linking surveys to ecological processes. Journal of Applied Ecology 52, 675–685. 10.1111/1365-2664.12432.

Douze M, Guzhva A, Deng C, Johnson J, Szilvasy G, Mazaré PE, Lomeli M, Hosseini L, Jégou H (2024). The FAISS library. 10.48550/arXiv.2401.08281.

Fabian Z, Miao Z, Li C, Zhang Y, Liu Z, Hernández A, Montes-Rojas A, Escucha R, Siabatto L, Link A, Arbeláez P, Dodhia R, Ferres JL (2023). Multimodal foundation models for zero-shot animal species recognition in camera trap images. 10.48550/arXiv.2311.01064.

Gabeff V, Rußwurm M, Tuia D, Mathis A (2024). WildCLIP: scene and animal attribute retrieval from camera trap data with domain-adapted vision-language models. International Journal of Computer Vision 132, 3770–3786. 10.1007/s11263-024-02026-6.

Gomez Villa A, Salazar A, Vargas F (2017). Towards automatic wild animal monitoring: identification of animal species in camera-trap images using very deep convolutional neural networks. Ecological Informatics 41, 24–32. 10.1016/j.ecoinf.2017.07.004.

Gretton A, Borgwardt KM, Rasch MJ, Schölkopf B, Smola A (2012). A kernel two-sample test. Journal of Machine Learning Research 13, 723–773. URL: https://jmlr.csail.mit.edu/papers/v13/gretton12a.html.

LILA BC (2024a). Snapshot Kgalagadi dataset. Labeled Information Library of Alexandria: Biology and Conservation. URL: https://lila.science/datasets/snapshot-kgalagadi.

LILA BC (2024b). Snapshot Kruger dataset. Labeled Information Library of Alexandria: Biology and Conservation. URL: https://lila.science/datasets/snapshot-kruger.

McNemar Q (1947). Note on the sampling error of the difference between correlated proportions or percentages. Psychometrika 12, 153–157. 10.1007/BF02295996.

Mulero-Pázmány M, Hurtado S, Barba-González C, Antequera-Gómez ML, Díaz-Ruiz F, Real R, Navas-Delgado I, Aldana-Montes JF (2025). Addressing significant challenges for animal detection in camera trap images: a novel deep learning-based approach. Scientific Reports 15, 16191. 10.1038/s41598-025-90249-z.

Nakagawa S, Lagisz M, Francis R, Tam J, Li X, Elphinstone A, Jordan NR, O’Brien JK, Pitcher BJ, Van Sluys M, Sowmya A, Kingsford RT (2023). Rapid literature mapping on the recent use of machine learning for wildlife imagery. Peer Community Journal 3, e35. 10.24072/pcjournal.261.

Norouzzadeh MS, Nguyen A, Kosmala M, Swanson A, Palmer MS, Packer C, Clune J (2018). Automatically identifying, counting, and describing wild animals in camera-trap images with deep learning. Proceedings of the National Academy of Sciences 115, E5716–E5725. 10.1073/pnas.1719367115.

Oquab M, Darcet T, Moutakanni T, Vo H, Szafraniec M, Khalidov V, Fernandez P, Haziza D, Massa F, El-Nouby A, Assran M, Ballas N, Galuba W, Howes R, Huang PY, Li SW, Misra I, Rabbat M, Sharma V, Bojanowski P (2023). DINOv2: learning robust visual features without supervision. 10.48550/arXiv.2304.07193.

Pardo LE, Bombaci SP, Huebner S, Somers MJ, Fritz H, Downs C, Guthmann A, Hetem RS, Keith M, le Roux A, Mgqatsa N, Packer C, Palmer MS, Parker DM, Peel M, Slotow R, Strauss WM, Swanepoel L, Tambling C, Tsie N, et al. (2021). Snapshot Safari: a large-scale collaborative to monitor Africa’s remarkable biodiversity. South African Journal of Science 117. 10.17159/sajs.2021/8134.

Peng Z, Dong L, Bao H, Ye Q, Wei F (2022). BEiT v2: masked image modeling with vector-quantized visual tokenizers. 10.48550/arXiv.2208.06366.

Radford A, Kim JW, Hallacy C, Ramesh A, Goh G, Agarwal S, Sastry G, Askell A, Mishkin P, Clark J, Krueger G, Sutskever I (2021). Learning transferable visual models from natural language supervision. In: Proceedings of the 38th International Conference on Machine Learning. Vol. 139. Proceedings of Machine Learning Research. PMLR, pp. 8748–8763. URL: https://proceedings.mlr.press/v139/radford21a.html.

Santamaria JD, Isaza C, Giraldo JH (2026). WildIng: a wildlife image invariant representation model for geographical domain shift. 10.48550/arXiv.2601.00993.

Schneider S, Greenberg S, Taylor GW, Kremer SC (2020). Three critical factors affecting automated image species recognition performance for camera traps. Ecology and Evolution 10, 3503–3517. 10.1002/ece3.6147.

Sokolova M, Lapalme G (2009). A systematic analysis of performance measures for classification tasks. Information Processing & Management 45, 427–437. 10.1016/j.ipm.2009.03.002.

Stevens S, Wu J, Thompson MJ, Campolongo EG, Song CH, Carlyn DE, Dong L, Dahdul WM, Stewart C, Berger-Wolf T, Chao WL, Su Y (2024). BioCLIP: A Vision Foundation Model for the Tree of Life. arXiv: 2311.18803 [cs.CV]. URL: https://arxiv.org/abs/2311.18803.

Swanson A, Kosmala M, Lintott C, Simpson R, Smith A, Packer C (2015). Snapshot Serengeti, high-frequency annotated camera trap images of 40 mammalian species in an African savanna. Scientific Data 2, 150026. 10.1038/sdata.2015.26.

Tabak MA, Norouzzadeh MS, Wolfson DW, Newton EJ, Boughton RK, Ivan JS, Humm JM, Bay R, Anderson EM, Crooks KR, Lewis JS, Carver S, Davis AJ, Yang L, Marshall B, Mosby CE, Boyce WM, Pomerleau G, Smith D, Vasi H, et al. (2020). Improving the accessibility and transferability of machine learning algorithms for identification of animals in camera trap images: MLWIC2. Ecology and Evolution 10, 10374–10383. 10.1002/ece3.6692.

Tabak MA, Norouzzadeh MS, Wolfson DW, Sweeney SJ, VerCauteren KC, Snow NP, Halseth JM, Di Salvo PA, Lewis JS, White MD, Teton B, Beasley JC, Schlichting PE, Boughton RK, Wight B, Newkirk ES, Ivan JS, Odell EA, Brook RK, Lukacs PM, et al. (2019). Machine learning to classify animal species in camera trap images: applications in ecology. Methods in Ecology and Evolution 10, 585–590. 10.1111/2041-210X.13120.

Tuia D, Kellenberger B, Beery S, Costelloe BR, Zuffi S, Risse B, Mathis A, Mathis MW, van Langevelde F, Burghardt T, Kays R, Klinck H, Wikelski M, Couzin ID, van Horn G, Crofoot MC, Stewart CV, Berger-Wolf T (2022). Perspectives in machine learning for wildlife conservation. Nature Communications 13, 792. 10.1038/s41467-022-27980-y.

Vélez J, McShea W, Shamon H, Castiblanco-Camacho PJ, Tabak MA, Chalmers C, Fergus P, Fieberg J (2023). An evaluation of platforms for processing camera-trap data using artificial intelligence. Methods in Ecology and Evolution 14, 459–477. 10.1111/2041-210X.14044.

Vyskočil J, Picek L (2025). Towards zero-shot camera trap image categorization. In: Computer Vision, ECCV 2024 Workshops. Vol. 15624. Lecture Notes in Computer Science. 10.1007/978-3-031-92387-6_3.

Whytock RC, Świeżewski J, Zwerts JA, Bara-Słupski T, Koumba Pambo AF, Rogala M, Bahaa-el-din L, Boekee K, Brittain S, Cardoso AW, Henschel P, Lehmann D, Orbell C, Abernethy KA (2021). Robust ecological analysis of camera trap data labelled by a machine learning model. Methods in Ecology and Evolution 12, 1080–1092. 10.1111/2041-210X.13576.

Willi M, Pitman RT, Cardoso AW, Locke C, Swanson A, Boyer A, Veldthuis M, Fortson L (2019). Identifying animal species in camera trap images using deep learning and citizen science. Methods in Ecology and Evolution 10, 80–91. 10.1111/2041-210X.13099.

Wilson EB (1927). Probable Inference, the Law of Succession, and Statistical Inference. Journal of the American Statistical Association 22, 209–212. 10.1080/01621459.1927.10502953.

Yu X, Wang J, Kays R, Jansen PA, Wang T, Huang T (2013). Automated identification of animal species in camera trap images. EURASIP Journal on Image and Video Processing 2013, 52. 10.1186/1687-5281-2013-52.

